# Probing Altered Receptor Specificities of Antigenically Drifting Human H3N2 Viruses by Chemoenzymatic Synthesis, NMR and Modeling

**DOI:** 10.1101/2023.04.05.535696

**Authors:** Luca Unione, Augustinus N.A. Ammerlaan, Gerlof P Bosman, Frederik Broszeit, Roosmarijn van der Woude, Yanyan Liu, Shengzhou Ma, Lin Liu, Tammo Diercks, Ana Ardá, Robert P. de Vries, Geert-Jan Boons

## Abstract

Prototypic receptors for human influenza viruses are cell surface *N*-glycans carrying α2,6-linked sialosides. Under immune pressure, A/H3N2 influenza viruses have emerged with altered receptor specificities that appear to recognize α2,6-linked sialosides presented on extended *N*-acetyl-lactosamine (LacNAc) moieties. Here, molecular recognition features of such drifted hemagglutinin’s (HAs) are examined by chemoenzymatic synthesis of complex *N*-glycans having ^13^C-labeled monosaccharides at strategic positions. The labeled glycans were employed in 2D STD-^1^H and ^13^C-HSQC NMR experiments to pinpoint which monosaccharides of the extended LacNAc chain engage with evolutionarily distinct HAs. The NMR data in combination with computational and mutagenesis studies demonstrate that mutations distal to the receptor binding domain of recent HAs have created an extended binding site that can directly interact with the extended LacNAc chain. A fluorine containing sialyl-LacNAc derivative is used as NMR probe to derive relative binding affinities and confirmed the contribution of the extended LacNAc chain for binding.

## Introduction

Spill-over of influenza A virus (IAV) from an animal reservoir into the human population has caused four pandemics in the past century^1^. These viruses became seasonal strains that evade neutralization induced by prior infections or immunization. Seasonal IAV cause significant disease burden, infecting 9–35 million individuals annually and causing 12–56,000 deaths^2^.

IAV expresses hemagglutinin (HA) and neuraminidase (NA) as two major envelope glycoproteins. HA binds to sialic acid (Neu5Ac) of glycoconjugates on host cells to initiate infection, whereas NA facilitates the release of progeny viruses from infected cells by cleaving sialosides. Human IAVs recognize sialosides α2,6-linked to galactoside (Gal), which are structures that are abundantly found in the upper respiratory tract of humans. On the other hand, HAs of ancestorial avian viruses exhibit a preference for α2,3-linked sialosides that are found in the duck enteric and chicken upper respiratory tract^3–5^.

Human IAVs have a remarkable ability to evolve and evade neutralization by antibodies elicited by prior infections or vaccination. This antigenic drift is mainly caused by mutations in the receptor binding site at the globular head of HA^6^. A/H3N2 viruses, which are the leading cause of severe seasonal influenza illness^7^, exhibit a particularly fast antigenic drift. Mutational changes due to antigenic pressure have resulted in altered glycan binding properties. These changes were first noticed by a lack of agglutination of fowl red blood cells. Furthermore, these viruses also propagate poorly in common laboratory hosts such as MDCK cells and eggs, indicating they require glycan receptors that are not expressed by these cells or hosts^8,9^. Glycan microarray and other binding studies have indicated these viruses have lost the ability to bind to simple α2,6-linked sialoglycans^10^ and instead bind to bi-antennary *N*-glycans that have on both arm extended *N*-acetyl lactosamine (LacNAc) moieties terminating in an α2,6-linked sialoside. Computational studies indicated a binding mode in which the two sialosides of the bi-antennary *N*-glycan bridge the binding sites of a trimeric HA protein resulting in high avidity of binding^11^.

We have constructed a glycan microarray populated with bi-antennary *N*-glycans that more closely resemble structures expressed in human respiratory tissue, including glycans with one or two sialic acid moieties linked to LacNAc chains of different length^12^. Compound **1** (Fig. 1) was established as the minimal epitope for early A/H3N2 viruses, whereas those that emerged after 2000 (*e.g*. NL2003) did not bind to **1** and required as minimal epitope a compound such as **2** with one of its arms presenting a tri-LacNAc moiety having a terminal α2,6-linked sialoside. Microarray data combined with sequence alignment, reported crystal structures, and MD simulations indicated an alternative binding model for recent A/H3N2 of the 3C.2 clade in which mutations remote from the receptor binding domain resulted in a rotation of the side chain of Tyr159, creating an extended binding site. Furthermore, a 225G/D mutation reoriented the poly-LacNAc chain of the compound allowing it to establish interactions with this extended binding site.

**Figure 1.**
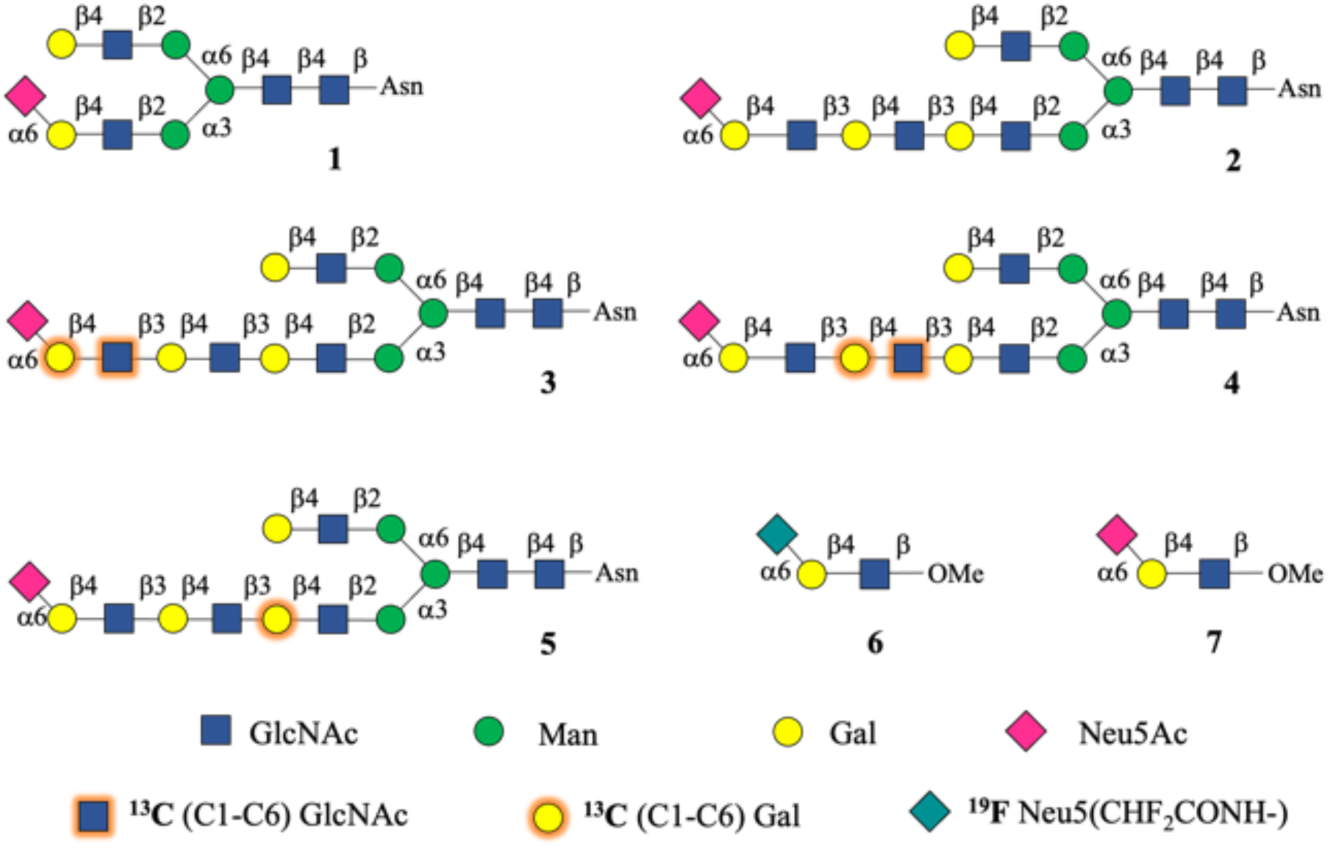
Isotopically glycans for NMR studies with evolutionary distinct HAs.

To experimentally validate this binding model, we describe here the chemo-enzymatic synthesis of *N*-glycan **2** and the ^13^C-labelled analogues **3**, **4** and **5** that differ in the positions of ^13^C-labeled monosaccharides along the poly-LacNAc chain. The ^13^C-labelled glycans were employed in 2D STD-^1^H,^13^C-HSQC experiments to pinpoint those monosaccharides of the poly-LacNAc chain of **2** that engage with HAs from evolutionarily distinct A/H3N2. The NMR data in combination with *in-silico* and mutagenesis studies demonstrated that mutations distal to the receptor binding domain of HA have created an extended binding site that can directly interact with the extended LacNAc chain of glycans. To investigate whether the affinity of HAs of contemporary A/H3N2 strains for the extended glycans has improved or compensated due to mutations at the receptor binding site, we employed a fluorine containing sialyl-LacNAc derivative (^19^F-SLN, compound **6** in Fig. 1) as a ^19^F NMR probe to derive relative binding affinities. This demonstrated that the additional contributions of the LacNAc chain improves binding affinity only for recent A/H3N2 strains.

The preparation of isotopomers **3**-**5** was critical for the STD-NMR experiments and uncover the contributions of each monosaccharide of the tri-LacNAc chain of **2** for binding to various HAs. This type of NMR experiment is a powerful approach to characterize glycans interactions with proteins at the atomic level^13^. Ligand protons in close proximity to the protein receive magnetic saturation resulting in strong STD signals. On the other hand, ligand protons that are more remote or do not engage with the protein exhibit weak or no STD signals. Large glycans suffer, however, from very low chemical shift dispersion of their ^1^H NMR signals precluding detailed binding studies by ^1^H-STD NMR. This is especially problematic when glycans present repeating units, such as poly-LacNAc chains, as these have almost indistinguishable ^1^H chemical shifts. These difficulties can be alleviated by using ^13^C isotope labels introduced at specific positions of an oligosaccharide chain. ^13^C filtered NMR spectra are then specific for these site selective probes and free of background signals from unlabelled monosaccharides within the *N*-glycan or from the HA glycoprotein. Such isotope labelling preserves binding characteristics while allowing heteronuclear NMR studies with atomic resolution and substantially increased sensitivity^14,15^. A challenge that we address in this study is the preparation of high complex glycan having selectively ^13^C labelled monosaccharides.

## Results and Discussion

### Synthesis of asymmetrical ^13^C labelled *N*-glycans

Glycan **2** and the three asymmetric isotopomers **3**-**5** (Fig. 1) were prepared from glycosyl asparagine **8** that could readily be prepared from a sialoglycopeptide extracted from egg-yolk powder followed by pronase treatment and then acid mediated hydrolysis of the sialosides (Scheme 1a)^16^. UDP-GlcNAc and UDP-Gal having ^13^C-isotopes at carbons 1 to 6 in combination with appropriate glycosyl transferases were employed to introduce ^13^C-labeled GlcNAc and Gal moieties into the target oligosaccharides. To selectively extend the α(1,3)-Man arm of **8** by an oligo-LacNAc chain, we exploited the inherent branch selectivity of β-galactoside α2,6-sialyltransferase 1 (ST6Gal1), which has a 10-fold higher activity for the α3-arm^17^, allowing for the convenient preparation of **9** (Scheme 1b,c). Next, the remaining terminal galactoside at the α6-arm of **9** was temporarily modified by an α1,2-fucoside by treatment with α1,2-fucosyltransferase 1 (FuT1) in the presence of GDP-Fuc to give **10**. The α1,2-fucoside temporarily blocks the α6-arm from modifications by mammalian glycosyl transferases^18^, and thus after removal of the α2,6-sialoside of **10** by acid mediated hydrolysis, the α3-arm of the resulting **11** could be selectively elaborated into a tri-LacNAc chain by the sequential use of B3GnT2 and B4GalT1 in combination with UDP-GlcNAc and UDP-Gal, respectively. The use of the ^13^C-labeled sugar donors for the installation of the first or second LacNAc moiety gave access to compounds **15** and **20** having the terminal and central LacNAc moiety labelled by ^13^C isotopes, respectively.

**Scheme 1.**
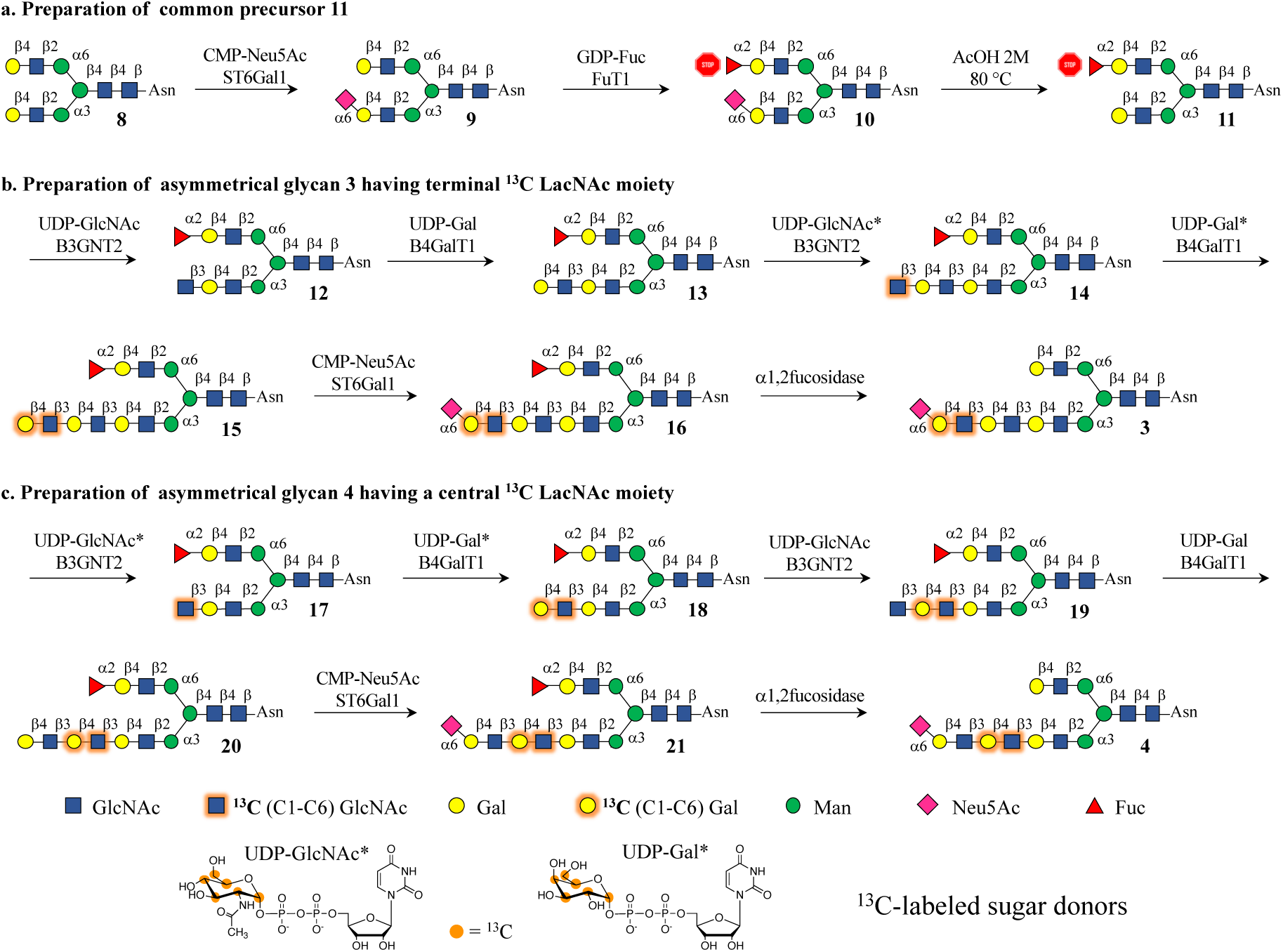
Chemoenzymatic synthesis of ^13^C labeled sialoglycans **3** and **4**. **a)** The α(1,3)-arm of glycan **8**, which was selectively modified as a 2,6-linked sialoside allowing selectively fucosylation of the α(1,6)-arm which temporarily block it glycosyl accepting properties (stop). After acid hydrolysis of the 2,6-linked sialoside, compound **11** was obtained that allows selective extension of the α(1,3)-arm. **b**) Glycan **3** was prepared by employing ^13^C labeled UDP-GlcNAc and UDP-Gal during the introduction of the final LacNAc assembly followed by treatment sialylation with ST6Gal1 and removal of the fucoside using a 1,2-fucosidase. **c**) Glycan **4** was prepared by using ^13^C labeled UDP-GlcNAc and UDP-Gal during the introduction of the central LacNAc moiety.

Target glycans **3** and **4** were obtained by treatment **15** and **20** with ST6Gal1 in the presence of CMP-Neu5Ac to give **16** and **21** that were treated with an αl,2-fucosidase to provide target compounds **3** and **4**.

To introduce the ^13^C labelled galactoside in the inner LacNAc moiety, symmetrical *N*-glycan **8** was treated with galactosyl hydrolase from *E. coli* that has high preference for the α3-arm^19^, to give after purification by HILIC-HPLC, asymmetrical glycan **22** (Scheme 2). The terminal galactoside at the α6-arm of **22** was temporarily blocked as α2,3-linked sialoside by treatment with β-galactoside α2,3-sialyltransferase 4 (ST3Gal4) in the presence of CMP-Neu5Ac to provide **23**. Next, a ^13^C-labelled galactoside was linked to the terminal GlcNAc moiety of **23** by employing B4GalT1 and ^13^C-labeled UDP-Gal to give **24**. The α3-arm of the latter compound was enzymatically extended into a tri-LacNAc structure to give **28** by repeated use of B3GnT2 and B4Gal1. Treatment of **28** with ST6Gal1 in the presence of CMP-Neu5Ac installed an α2,6-sialoside thereby providing **29**. Finally, the α2,3-linked sialoside at the α6-arm of **29** was selectively removed by treatment with α2,3-neuraminidase to afford target glycan **5**. The same synthetic strategy was used to synthetize unlabelled **2** from unlabelled UDP-sugar donor.

**Scheme 2.**
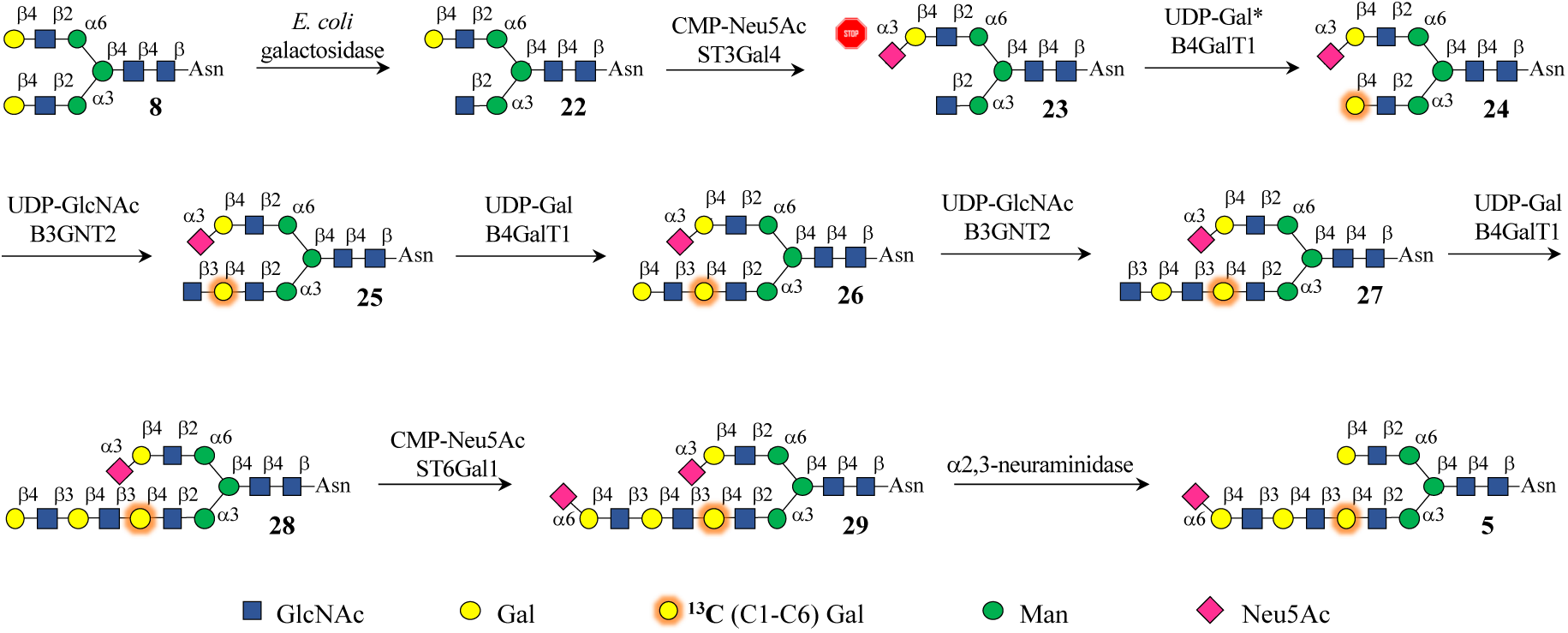
Chemoenzymatic synthesis of glycan **5**. Selective removal of the galactoside of the α3-arm allowed the selective installation of the ^13^C-labeled galactoside at the first residue of the tri-LacNAc moiety to give entry into the synthesis of glycan **5**.

### 1D ^1^H-STD NMR experiments

HAs from evolutionarily different time points, HK68 (1968), NL91 (1991) and NL03 (2003), were selected to analyse their binding preferences for unlabelled glycan **2** by 1D ^1^H-STD NMR experiments (Fig. 2a). HK68 is the prototypical HA of the 1968 Hong-Kong pandemic that was adapted from its avian predecessor to bind human-type receptors^20^. NL91 HA was isolated when these viruses could still hemagglutinate fowl erythrocytes. Later in the antigenic evolution, these viruses lost this property as was observed for in the NL03 strain. The STD signals confirmed that all three HAs have functional binding properties. As expected, severe ^1^H chemical shift overlap limited ^1^H-STD based analysis that nonetheless revealed clear differences. Thus, HA from HK68 showed STD signals only for the sialic acid moiety, indicating that only this residue contributes to protein binding. The strongest STD signals were observed for H4, H7, H8, H9 and the methyl group of the acetamide moiety of the sialoside. In contrast, the ^1^H-STD spectrum of NL91 HA showed several additional STD signals, including the methyl group of an acetamide of a GlcNAc residue. Finally, the ^1^H-STD spectrum of NL03 HA was substantially more complex and showed, among others, STD signals above 4 ppm that correspond to the H1 and H4 of galactosides. It was, however, not possible to identify specific monosaccharides due to the chemical shift degeneration of the nuclei of the tri-LacNAc chain. These results indicate that from 1968 to 2003, A/H3N2 HAs evolved to recognize additional residues of an extended LacNAc chain beyond the terminal α2,6-sialoside.

**Figure 2.**
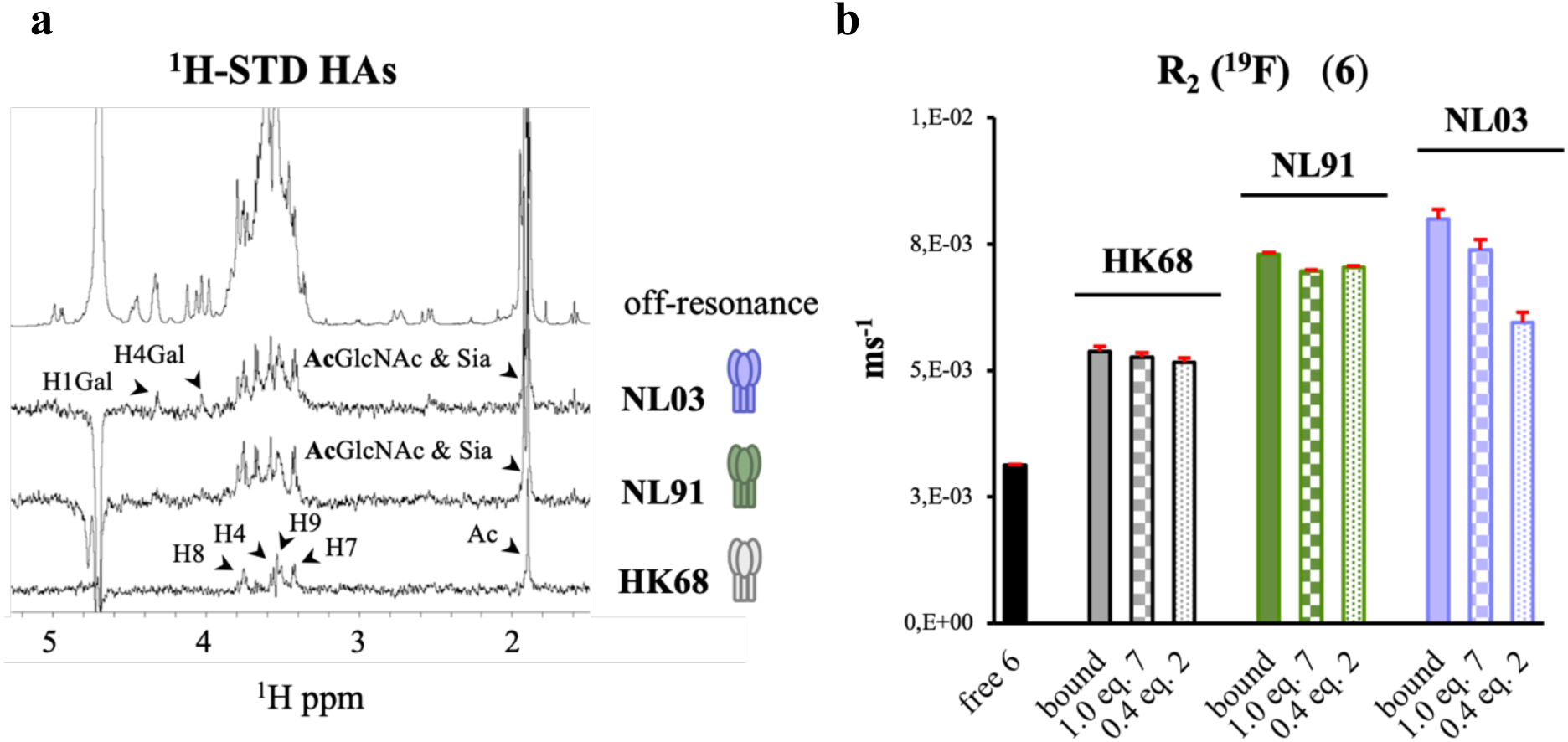
NMR binding studies. a) 1D ^1^H-STD NMR. Spectra obtained for the complexes of HK68, NL91, and NL03 HAs with the glycan **2**. b) ^19^F-R2 NMR. Comparison of transverse ^19^F relaxation rate, R_2_(^19^F), for the fluorinated probe (**6**) in the absence or presence of the different HA proteins. Addition of competitors molecules **7** and **2** causes a decrease in R_2_(^19^F) proportional to their affinities relative to the probe compound **6**.

### Transverse relaxation rate R_2_(^19^F) NMR experiments to establish relative binding affinities

To identify the minimal binding epitope and derive relative binding affinities for the three HAs, we used fluorine containing Neu5DFA(α2,6)LacNAc (^19^F-SLN, compound **6**, Fig. 1) as probe (see supplementary section 2 for details of the synthesis of **6**). The strategic incorporation of a non-endogenous fluorine atom (^19^F) enables ligand-protein interaction studies by molecule specific ^19^F NMR experiments by monitoring R_2_(^19^F)-relaxation rates that do not suffer from overlap or dispersion problems as in ^1^H-based NMR experiments described above^21^. First, the overall transverse relaxation rate was measured, R_2_(^19^F), of ^19^F-SLN in the absence of HA. Addition of the HAs to this fluorinated probe caused a large increase in R_2_(^19^F) relative to the free form indicating ^19^F-SLN binding (Fig. 2b, filled bars). R_2_(^19^F) increases for NL91 and NL03 were substantially larger than for HK68. These observations indicate a substantial affinity increase for the ^19^F-SLN trisaccharide from HK68 to NL91 and slightly increased from NL91 to NL03 as previously also observed by ELISA assays^11^. Together with the results from the 1D ^1^H-STD NMR experiments, it is concluded that the higher affinity of NL91 and NL03 HAs for this trisaccharide is due to additional recognition of the Gal and GlcNAc residues absent in HK68.

As the replacement of two hydrogen atoms by fluorines at the acetamide moiety may alter ligand binding properties^22^, the suitability of compound **6** as a probe was examined by a competition experiment using unlabelled sialyl LacNAc **7** (Fig. 1). Thus, 1.0 equivalent of **7** was added to the NMR tube containing the HA proteins and ^19^F-SLN. Addition of the nonfluorinated competitor caused a decrease in the overall R_2_(^19^F) relaxation rate of the fluorinated probe (Fig. 2b, square-pattern filled bars) confirming its functional binding at the same site of the receptor HAs. Next, we examined the relative HA affinities for compound **2** that present an α2,6-linked sialoside on its extended tri-LacNAc chain. Thus, 0.4 equivalent of **2** was added relative to the fluorinated probe to samples containing the three different HA proteins and ^19^F-SLN (**6**), (Fig. 2b, dotted-pattern filled bars). For the HA’s from HK68 and NL91, the addition of the competing glycan **2** caused a reduction of R_2_(^19^F) for ^19^F-SLN similar to that induced by trisaccharide **7**, indicating that the two additional LacNAc moieties in **2** do not enhance the binding affinity. On the other hand, for the HA of NL03 a substantially stronger reduction of R_2_(^19^F) was observed indicating that the inner di-LacNAc moiety contribute to recognition by providing additional intermolecular interactions in agreement with the ^1^H-STD result.

### 2D ^1^H-STD-^1^H,^13^C-HSQC experiments

To elucidate specific contributions of each monosaccharide of the poly-LacNAc chain of **2** to HA binding, we recorded 2D ^1^H-STD-^1^H,^13^C-HSQC NMR experiments for isotopomers **3**, **4**, and **5** in the presence of the three HAs.

This variant of the ^1^H-STD experiment covers ^1^H[^13^C]-STD effects only for protons bound to a ^13^C isotope and, therefore unambiguously demonstrating whether specifical ^13^C labelled monosaccharides contribute to recognition. ^1^H-STD-^1^H,^13^C-HSQC experiments with isotopomer **3** showed clear differences in recognition between HK68, NL91, and NL03 HA (Fig. 3a). For HK68, no ^1^H[^13^C]-STD signals were observed indicating that the terminal LacNAc unit is not part of the recognised epitope. In contrast, ^1^H[^13^C]-STD signals were observed with moderate and strong intensities for NL91 HA and NL03 HA, respectively, indicating that these HAs evolved to recognise sialic acid linked to a LacNAc moiety.

**Figure 3.**
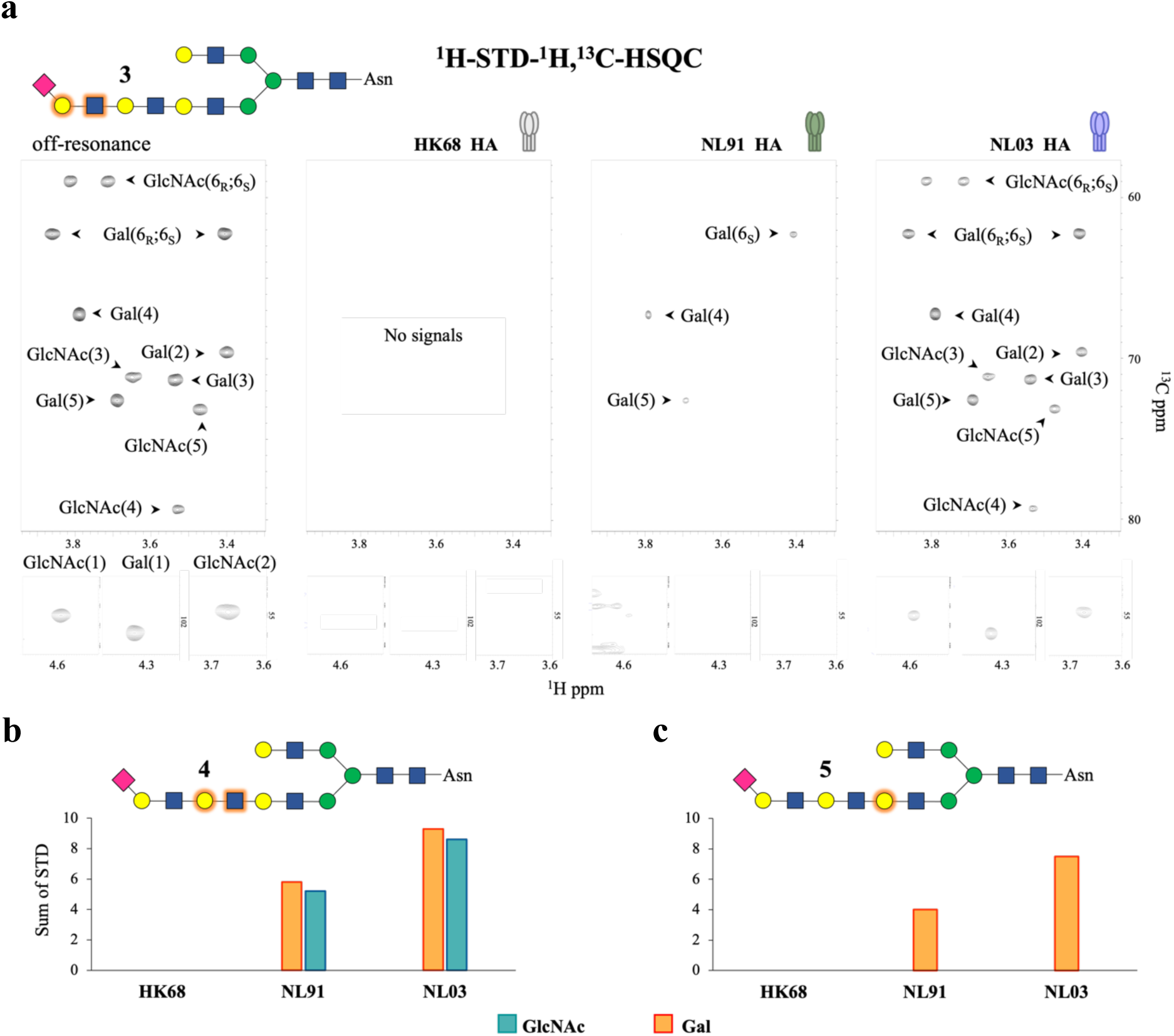
2D ^1^H-STD-^1^H,^13^C-HSQC NMR experiments probing the interaction of compounds **3** to **5** with different HA proteins. a) Off-resonance reference (left) and associated ^1^H-STD-^1^H,^13^C-HSQC spectra for the indicated three HA proteins in complex with compound **3**. b, c) Sum of all ^1^H[^13^C]-STD signals of the ^13^C labelled Gal (orange) and GlcNAc (cyan) moieties in compounds **4** (b) and **5** (c) in complex with the three HA proteins.

Similar experiments with isotopomer **4**, in which the central LacNAc unit is ^13^C-labelled uncovered further differences in recognition by the three HAs (Fig. 3b and Supplementary Fig. 4). As expected, HK68 HA did not produce any ^1^H[^13^C]-STD signal while the HA from NL91 and NL03 clearly invoked signals. The ^1^H[^13^C] STD signals are derived from both Gal and GlcNAc and were stronger for NL03 HA, where the sum of all STD signal was almost twice as those observed for NL91 HA, indicating that the central LacNAc moiety is more intimately bound by NL03. Finally, ^1^H-STD-^1^H,^13^C-HSQC experiments with isotopomer **5** (Fig. 3c and Supplementary Fig. 5), which has a ^13^C-labeled Gal farthest away from the sialoside, did not show any ^1^H[^13^C]-STD signals for HK68 HA. Thus, this most ancient viral HA recognises the glycan exclusively through the terminal α2,6-linked sialoside. NL91 HA produced weak ^1^H[^13^C]-STD signals while NL03 HA gave rise to the strongest STD signals, where the sum of ^1^H[^13^C]-STD intensities was almost twice as those observed for NL91. The most intense ^1^H[^13^C]-STD signals from **5** in complex with NL03 HA corresponded to the H3, H4 and H5 protons of the ^13^C labelled Gal residue (Supplementary Fig. 6). In galacto-configurated monosaccharides, these protons point in the same direction, forming a so-called alpha-face of the pyranoside rings, and often participate in CH-π interactions with aromatic amino acids receptor proteins^23^.

The 2D ^1^H-STD-^1^H,^13^C-HSQC experiment in combination with residue specific ^13^C labelling of glycan **2** made it possible to disentangle the contributions of individual monosaccharides of the extended poly-LacNAc chain for recognition by three different HAs. Increasing ^1^H[^13^C]-STD signal intensities were observed during evolution of HA evolution indicating more intimate contacts with the poly-LacNAc chain. Thus, for the original pandemic A/H3N2 human virus (HK68) the poly-LacNAc moiety does not contribute to binding, whereas, for later strains, site-specific mutations close to its glycan binding site enable further interactions with the extended glycan chain. For the evolutionarily early NL91 HA, these interactions are mostly limited to the sialic acid-linked galactoside while the more contemporary NL03 HA recognises almost the entire poly-LacNAc chain.

### Modelling of glycan-HA complexes

To uncover the binding modes of the three HAs, we combined the results from the NMR experiments with *in silico* modelling (Fig. 4). Full atom 1 μs molecular dynamic (MD) simulations in explicit water were performed for asymmetric *N*-glycan **2** in complex with the three HAs. In agreement with previous X-ray studies^24^, the complex of **2** with HK68 HA showed that the recognition of sialic acid is mediated by hydrophobic interactions with amino acids Y98, H183, W153 and L226, as well as by hydrogen bonding interactions with G135-N137, S228, and E190 (Supplementary Fig. 7a). Analysis of the MD trajectory showed that E190 engages the Sia-1 OH9 through hydrogen bonding with an average distance of ~2.9 Å, while L226 makes hydrophobic interactions with the sialic acid-galactose glycosidic linkage. Beyond the sialic acid residue, the MD simulation did not reveal stable intermolecular interactions and throughout the MD trajectory the poly-LacNAc chain was flexible and mainly solvent exposed (Supplementary Fig. 8).

**Figure 4.**
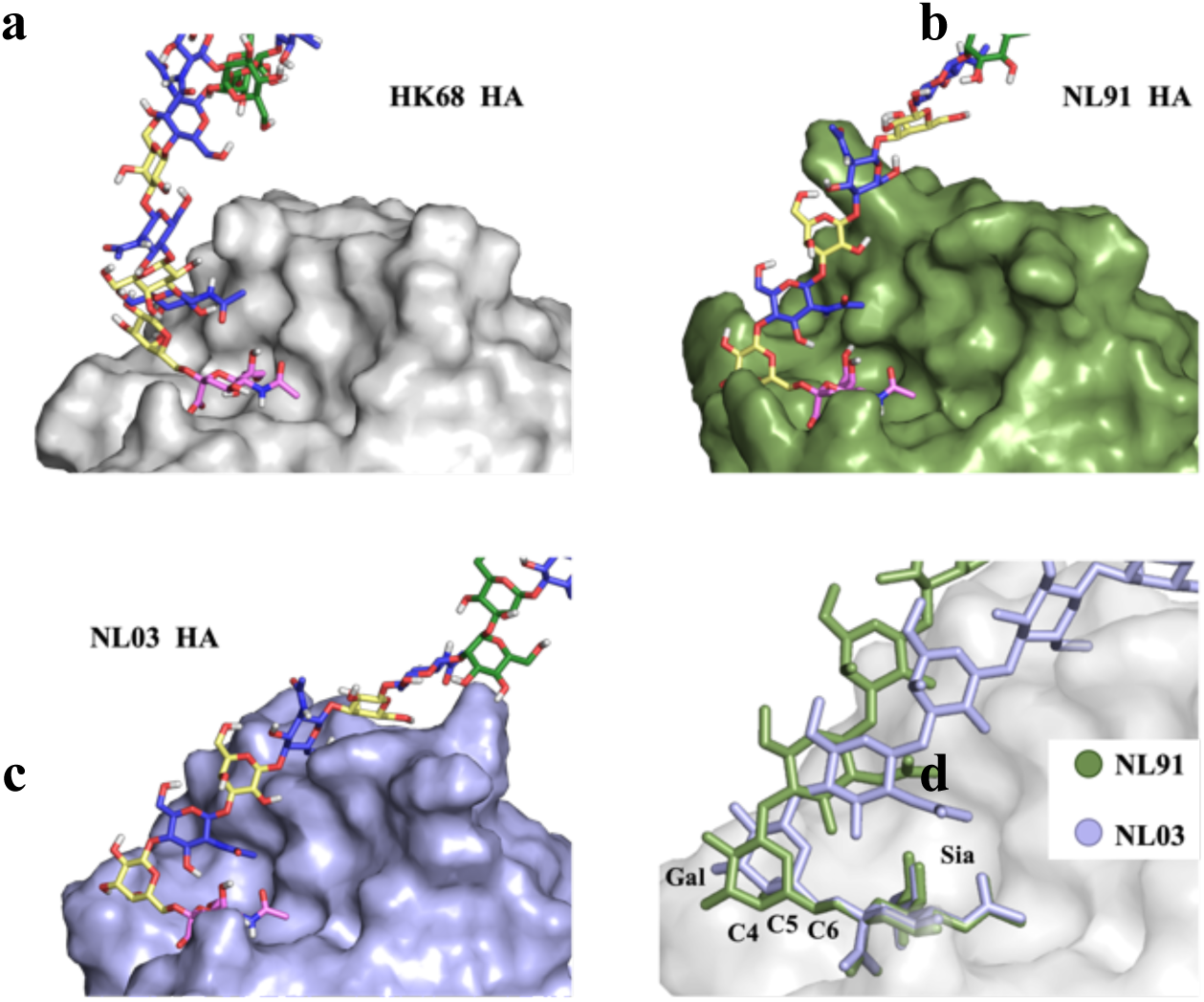
Models of *N*-glycan **2** binding by different HA proteins. a) Binding of **2** by HK68 HA. Only the terminal sialic acid fits into the protein binding pocket while the rest of the poly-LacNAc chain is solvent exposed. b) Binding of **2** by NL91 HA. The sialic acid and rearby parts of the α2,6-linked galactoside establish contacts with the protein. c) Binding of **2** by NL03 HA. The entire poly-LacNAc chain establishes contacts with the protein. d) Comparison of glycan binding modes for NL91 and NL03 HA. A change in the C5-C6 torsion angle adjacent to the Neu5Ac-Gal glycosidic linkage facilitates a reorientation of the underling glycan chain for extended interaction with NL03 HA.

The MD simulation for NL91 HA (Supplementary Fig. 7b and 9) revealed that the S145K mutation, at the sialic acid binding site, enabled a new hydrogen bond compensating for loss of a hydrogen bond to the carboxylic acid of Neu5Ac from a N137Y mutation. The new Y137 side chain in NL91 HA is properly oriented to face the sialic acid-galactose glycosidic bond, which is consistent with the STD signals detected for H4 to H6_R/S_ of this Gal residue (Fig. 3a) that were unobservable for HK68 HA. Another relevant NL91 HA mutation is Q189R in the 190 helix where the more extended side chain of the R residue is oriented towards solvent. This leaves space for the acetamide group of the GlcNAc (GlcNAc-3) to be accomodated into a hydrophobic pocket of the protein, where its methyl group makes Van der Waals contacts with L194. These results are consistent with the observed ^1^H-STD signal for the acetamide moiety of the GlcNAc residue in NL91 HA (Fig 2a).

The MD trajectory for NL03 HA in complex with **2** (Supplementary Fig. 7c and 10) showed that the further Y137S mutation restores a hydrogen bond with the carboxylic acid of Neu5Ac as originally observed for HK68, while maintaining the K145 hydrogen bond to OH4 of the sialoside, as in NL91. Yet, the concomitant E190D mutation now precludes hydrogen bonding with OH9 of the sialoside due to the shorter D side chain. The G225D mutation appears crucial to switch the glycan orientation in the receptor binding site consistent with the observed stronger ^1^H-STD signal for the terminal Gal in NL03 relative to NL91 HA. The new hydrogen bonding between D225 side chain and both OH4 and OH3 in Gal forces the α2,6-sialyl-Gal glycosidic linkage in a conformation that moves the underlying glycan chain towards the HA 190-helix (Fig. 4d). In this orientation, the hydrophobic interaction between GlcNAc-3 acetamide moiety and L194 is stabilized because S193 can establish an additional hydrogen bond with Gal-4. Further mutations distal to the RBS in NL03 HA reorient the Y159 side chain to face the Gal-6 and establish CH-π interactions (Supplementary Fig. 7c). The MD results are in accordance with the NMR data showing a substantial increase in ^1^H-STD signal intensities for H3, H4, and H5 of the Gal-6 residue. Thus, four residues (D225, L194, S193, and Y159) in the binding pocket of NL03 HA accommodate the extended glycan chain and contribute to the strong ^1^H[^13^C]-STD signals observed for compounds **3**, **4**, and **5** in the presence of NL03 HA.

### Structural studies using additional variants and mutagenesis to confirm the binding models

We expanded our study by including two additional HAs having further mutations in the extended binding site. NL09 HA (2009) has K145N, Y159F, S193F, and D225N mutations that according to our models are expected to impair recognition of glycan **2**. Indeed, the 2D ^1^H-STD-^1^H,^13^C-HSQC experiment with the three isotopomers **3**, **4**, and **5** showed substantially weaker ^1^H[^13^C]-STD signals than observed for NL03 HA (Fig. 5a). While MD simulation revealed that N145 cannot engage the sialic acid by hydrogen bonding interaction in the way K145 does for NL03 HA (Supplementary Fig. 7d). Furthermore, the substitution of D by N at residue 225 results in a weaker hydrogen bond acceptor while the S193F mutation precludes hydrogen bond interaction with the Gal in the central LacNAc unit. Finally, the Y159F mutation produces a weaker CH-π acceptor^25^.

**Figure 5.**
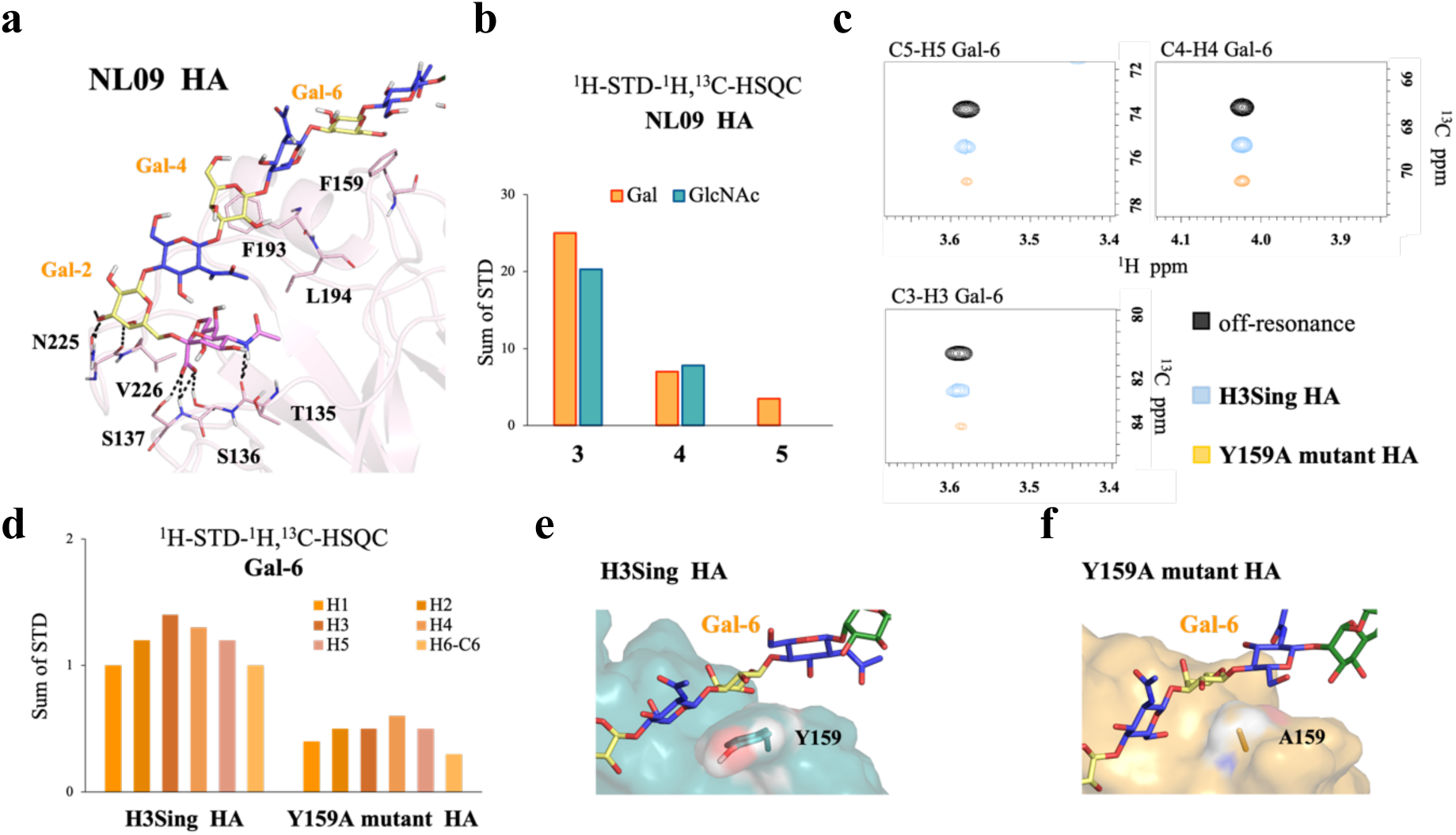
Glycan binding by additional HA variants. a-b) Glycan binding by NL09 HA. MD derived complex structure with **2** (a), and ^1^H[^13^C]-STD intensities from the 2D ^1^H-STD-^1^H,^13^C-HSQC spectrum of **3**-**5**. c) Comparison of ^1^H[^13^C]-STD signals of **5** in the presence of H3Sing HA and its Y159A HA mutant. Note the significantly weaker STD signals for mutant H3SingY159A HA relative to the wild type. d) ^1^H-STD intensities for the ^13^C-labelled galactoside residue in **5**. e-f) Model of glycan-HA interactions in H3Sing HA (e) and its Y159A HA mutant (f). The absence of an aromatic moiety in the Y159A mutant precludes the key CH-π interaction with Gal6.

Mutational reversibility has been observed in HAs^26^, and for example in H3Sing HA (2016) residues 225 and 159 reverted back to D and Y, respectively, as in NL03 HA, and residue 193 back to F, as in NL09 HA. MD simulations confirmed that the key hydrogen bond between D225 and Gal-2 has been regained (Supplementary Fig. 7e) while the bulky side chain of F193 provokes steric clashes with the glycan chain as in NL09 HA. Finally, Y159 re-established strong CH-π interactions with Gal-6. Accordingly, NMR experiments showed strong ^1^H[^13^C]-STD signals for the H3, H4, and H5 in Gal-6 as observed for NL03 HA.

To further assess the key contribution of Y159 for receptor binding, we produced a mutant of the H3Sing HA variant in which this tyrosine moiety is replaced by alanine, thereby abrogating the CH-π interaction. The NMR experiments showed substantial weaker ^1^H[^13^C]-STD signals for Gal-6 (Fig. 5c-d) indicating much less intimate contacts. These results underpin that several residues cooperate in glycan binding.

## Conclusion

Human influenza A viruses have a remarkable ability to evolve and evade neutralization by antibodies elicited by prior infections or vaccination. This antigenic drift is mainly caused by mutations of amino acid in the globular head of HA where binding occurs with sialic acid containing receptors of host cells. As a result, receptor-binding modes of IAVs co-evolve usually by orchestrated mutations of several amino acids that allow immune evasion but maintain glycan binding capabilities.^24^ A/H3N2 viruses, which are the leading cause of severe seasonal influenza illness^7^, exhibit a particularly fast antigenic drift and have evolved to recognize sialosides presented on extended LacNAc moieties. Predicting influenza evolution requires an understanding, at a molecular level, how mutational changes driven by immune pressure shape glycan recognition. X-ray crystallography has provided a key understanding of the recognition of sialosides by HAs, however, such an approach is difficult to implement to examine recognition of complex glycans such as **2** because of difficulties of forming co-complexes.

STD-NMR is an attractive solution-based approach that can reveal which protons of a ligand interaction with a protein and in combination with computational approaches can provide models of glycan-protein complexes. The implementation of such an approach for complex glycans is, however, complicated by NMR proton degeneracy. In this study, we addressed this problem by chemically synthesizing complex N-glycans having at specific positions ^13^C labeled monosaccharides. To obtain the targeted isotopomers, we employed an N-glycan isolated for a natural source as starting material, recombinant human glycosyl transferases and ^13^C labeled sugar nucleotide donors. Arm-selective extension of one of the arms of a symmetrical *N*-glycan starting material was possible by exploiting inherent branch selectivities of glycosyl transferases and glycosidases combined with modifications that temporarily block one of the arms from further extension. The use of the synthetic isotopomers in 2D ^1^H-STD-^1^H,^13^C-HSQC experiments made it possible to unravel the contribution of individual monosaccharides of an extended poly-LacNAc chain of an N-glycan for binding to evolutionary distinct HAs. In the case of the HA of the original pandemic A/H3N2 (HK68), only the sialoside engages with the protein and no contribution of the poly-LacNAc chain for binding was detected. For later strains, mutations close to the glycan binding site enabled interactions with the LacNAc chain. For the HA of the evolutionarily early NL91, these interactions are mostly limited to the sialic acid-linked galactoside while the more contemporary NL03 HA recognised almost the entire poly-LacNAc chain. The latter was confirmed by a fluorine containing sialyl-LacNAc derivative as NMR probe that provided relative binding affinities and demonstrated additional contributions of the extended

LacNAc chain for binding. Combination of the NMR data with molecular modelling revealed how specific residues along the extended binding site of HA synergistically operate in glycan recognition. In particular, a G225D mutation changes the glycan orientation and moves the LacNAc chain closer to the protein surface. A distal mutation to the receptor binding site in NL03 HA reorient the Y159 side chain to face the Gal-6 making it possible to establish CH-π interactions. Mutagenesis experiments confirmed the important of the Y159 side chain for interacting with the extended LacNAc chain. We expect that the generated structural models of different HA-glycan complexes will facilitate the development of predictive models for the evolution of IAV variants.

## Methods

### Enzymatic Synthesis

*Materials and general protocols*. Glycosyltransferase were expressed as previously described (St6Gal1, Fut1, St3Gal4, B4Galt1 and B3Gnt2)^27^. Galactosidase of *E. coli* was purchased from Sigma Aldrich [Cat# G5635] α1,2-Fucosidase [Cat# P0724S] and α2,3-Neuraminidase [Cat# P0743] were purchased from New England BioLab. Alkaline phosphatase (FastAP) was purchased from Thermo Scientific [Cat# EF0651]. Uridine 5’-diphosphogalactose (UDP-Gal), uridine 5’-diphospho-N-acetyl-glucosamine (UDP-GlcNAc) and cytidine-5’-monophospho-N-acetylneuraminic acid (CMP-Neu5Ac) were obtained from Roche Diagnostics [UDP-Gal: Cat# 07703562103; UDP-GlcNAc: Cat# 06369855103; CMP-Neu5Ac: Cat# 05974003103]. ^13^C labelled Uridine 5’-diphosphogalactose (UDP-Gal) and uridine 5’-diphospho-N-acetyl-glucosamine (UDP-GlcNAc) were obtained from Omicron [UDP-GlcNAc C1-C6: Cat# NTS-009; UDP-Gal C1-C6: Cat# NTS-005]. Reaction mixtures were purified by a size exclusion Biogel (P2) from BioRad in Econo glass columns (0.7 x 30 cm / 1.5 x 30 cm / 1.5 x 50 cm) coupled to a BioFrac fraction collector (BioRad). Carbohydrate-containing fractions were detected by thin layer chromatography and an appropriate staining reagent (15 mL AcOH and 3.5 mL p-anisaldehyde in 350 mL EtOH and 50 mL H_2_SO_4_). Final products were purified by high performance liquid chromatography (HPLC) using an XBridge HILIC column (10 mm (0) x 250 mm (l), 5 μm particle size) on a semi-preparative liquid chromatography system from Shimadzu (LC-20AT, SIL-20A, CBM-20A, SPD-20A, FRC-10A). The purification was performed using 10 mM NH_4_HCO_3_ in 10% H_2_O in MeCN (buffer B) and 10 mM NH_4_HCO_3_ in 100% H_2_O (buffer A). The progress of the reactions was monitored on a Shimadzu liquid chromatography mass spectrometry (LC-MS) (system controller: SCL10A-VP; HPLC pumps: LC10AD-VP; injector: SIL10AD-VP) using a ZIC HILIC column (ZeQuant, PEEK coated guard HPLC column, 3.5 μm particle size, 20 x 2.1 mm). The LC system was attached to a Bruker Daltonics microTOF-Q mass spectrometer.

### General procedures for synthesis

#### Acid mediated hydrolysis of Neu5Ac

The substrate was dissolved in an aqueous solution of acetic acid (2 M) and kept at 65 °C for 24 h. The solvents were removed under an N2 flow and the resulting mixture was applied to size exclusion chromatography. Carbohydrate-containing fractions were lyophilized and used without further purification.

#### Installation of α2,6-linked Neu5Ac using ST6Gal1

Acceptor and CMP-Neu5Ac (1.5 eq) were dissolved in a Tris buffer (50 mM, pH 7.3, 0.1 wt% BSA) to obtain an acceptor concentration of 2 mM. ST6Gal1 (42 μg per μmol acceptor) was added to the mixture and the resulting reaction mixtures was incubated overnight at 37 °C with gentle shaking. The progress of the reaction was monitored by LC-MS. In case of incomplete conversion, additional ST6Gal1 (20 μg per μmol acceptor) was added and the reaction mixture was incubated at 37 °C for an additional 24 h. After completion of the reaction, the enzyme was removed by spin filtration using a 10 kDa cut-off filter, the filtrate was lyophilized and applied to size exclusion chromatography. Carbohydrate-containing fractions were collected, concentrated by freeze drying and either used for further modification or further purification by HPLC.

#### Installation of a1,2-linkedfucoside using Fut1

Acceptor and GDP-Fuc (1.3 eq) were dissolved in a Tris buffer (50 mM, pH 7.3, 0.1 wt% BSA) containing MnCl2 (10 mM) to a final acceptor concentration of 5 mM. FuT1 (7 μg per μmol acceptor) were added and the reaction mixture was incubated at 37 °C for 72 h with gentle shaking. The progress of the reaction was monitored by LC-MS.

#### Installation of β1,3-linked GlcNAc using B3GnT2

Acceptor and UDP-GlcNAc (1.5 eq) were dissolved in a HEPES buffer (50 mM, pH 9.6, 0.1 wt% BSA) containing DTT (1 mM) and MnCl2 (20 mM) to obtain a concentration of 5 mM. B3GnT2 (30 μg per μmol acceptor) and CIAP (1 u μL-1, 1 u per μmol of added nucleotide) were added to the mixture and then incubated overnight at 37 °C with gentle shaking. The progress of the reaction was monitored by LC-MS. In case of incomplete conversion, additional UDP-GlcNAc (0.5 eq), CIAP (1 u μL-1, 1 u per μmol of added nucleotide) and B3GnT2 (15 μg per μmol acceptor) were added and the reaction mixture incubated at 37 °C for an additional 24 h. The reaction mixture was lyophilized and applied to size exclusion chromatography. Carbohydrate-containing fractions were lyophilized and used without further purification.

#### Installation of β1,4-linked galactoside using B4GalT1

Acceptor and UDP-Gal (1.5 eq) were dissolved in a Tris buffer (50 mM, pH 7.3, 0.1wt% BSA) containing MnCl2 (20 mM) to obtain a concentration of 5 mM. B4GalT1 (20 μg per μmol acceptor) and CIAP (1 u μL-1, 1 u per μmol of added nucleotide) were added and the resulting reaction mixture was incubated for 18 h at 37 °C with gentle shaking. The progress of the reaction was monitored by LC-MS. In case of incomplete conversion, additional UDP-Gal (0.5 eq), CIAP (1 u μL-1, 1 u per μmol of added nucleotide) and B4GalT1 (10 μg per μmol acceptor) were added and the reaction mixture incubated at 37 °C for a further 24 h. The reaction mixture was lyophilized and applied to size exclusion chromatography. Carbohydrate-containing fractions were lyophilized.

#### Fucoside hydrolysis

The fucosylated glycan was dissolved into 50 mM sodium acetate buffer at pH 5.5, 5mM CaCl2, and treated with the broad specific α1-2,3,4,6 fucosidase (80 units). The reaction mixture was incubated at 37 °C with gentle shaking. The progress of the reaction was monitored by LC-MS.

#### Galactoside hydrolysis

The galacto-containing glycan was dissolved in 50 mM TRIS buffer at pH 7.3, 15 mM MgCl2 and treated with galactosidase (20 units). The reaction mixture was incubated at 37 °C with gentle shaking. The progress of the reaction was monitored by LC-MS.

#### Installation of α2,3-linked Neu5Ac using ST3Gal4

Acceptor and CMP-Neu5Ac (1.5 eq) were dissolved in a Tris buffer 50 mM, pH 7.5, 1% v/v BSA (from stock solution 1 mg/mL) and 1% v/v CIAP (from stock solution 20U/μL) to a final acceptor concentration of 5 mM. ST3Gal4 (42 μg per μmol acceptor) was added and the reaction mixture was incubated overnight at 37 °C with gentle shaking. The progress of the reaction was monitored by LC-MS.

#### Enzymatic hydrolysis of α2,3-linked Neu5Ac

The α2,3-sialylated glycan was dissolved into 50mM sodium acetate buffer at pH 5.5, 5mM CaCl2, and treated with the high specific α2,3-Neuraminidase S (160000 U/mg). The reaction mixture was incubated at 37 °C with gentle shaking. The progress of the reaction was monitored by LC-MS.

### HA expression

Recombinant trimeric IAV hemagglutinin proteins open reading frames were cloned into the pCD5 expression vector as described previously^28^, in frame with a GCN4 trimerization motif (KQIEDKIEEIESKQKKIENEIARIKK), a superfolder GFP^29^ and the Twin-Strep-tag (WSHPQFEKGGGSGGGSWSHPQFEK); IBA, Germany). The open reading frames of the HAs of A/Hong-Kong/1/1968 H3 (AFG71887.1), A/Netherlands/816/1991 H3 (EPI_ISL_114608), A/Netherlands/109/2003 H3 (EPI_ISL_113016), A/Netherlands/761/2009 H3 (EPI_ISL_1107270), A/Singapore/INFH-16-0019/2016 H3 (3C.2a) (QQY97257.1), were synthesized and codon optimized by GenScript.

The trimeric HAs were expressed in HEK293T cells with polyethyleneimine I (PEI) in a 1:8 ratio (μg DNA:μg PEI) for the HAs as previously described^28^. The transfection mix was replaced after 6 hours by 293 SFM II suspension medium (Invitrogen, 11686029), supplemented with sodium bicarbonate (3.7 g/L), Primatone RL-UF (3.0 g/L, Kerry, NY, USA), glucose (2.0 g/L), glutaMAX (1%, Gibco), valproic acid (0.4 g/L) and DMSO (1.5%). Culture supernatants were harvested 5 days post transfection and purified with sepharose strep-tactin beads (IBA Life Sciences, Germany) according to the manufacturer’s instructions.

### NMR sample preparation and analysis

Proteins and ligands were dissolved in a deuterated TRIS-d11 50mM buffer pD 7.8 in D2O. Shigemi NMR tubes with a diameter of 3 mm were used. ^1^H-STD and ^1^H,^13^C-STD HSQC experiments were acquired with a HA proteinsconcentration of 5 μM and a ligand at 300 μM (ratio 1:60). STD-experiments were acquired on a 800 MHz BRUKER AVANCE III spectrometer, equipped with a TCI cryo-probe with z-gradient coil and TopSpin 3.2.7 (BRUKER) software was employed for data acquisition and processing. The temperature was set to 293 K. The 2D STD ^1^H,^13^C-HSQC experiment was previously reported^14^. Briefly, the ^1^H,^1^H-STD module was implemented with saturation by a train of 4 x 90° PC9_4 shaped pulses (with 1 ms separation) during d9 = 2 s, applied at the methyl ^1^H peak (0.84 ppm) with the non-saturation cut-off set at 1.18 ppm (i.e. distance Δν = 0.34 ppm), resulting in a PC9_4 pulse length of 33 ms at 800 MHz field strength. Protein saturation was alternated with off-resonant irradiation (at −25 ppm) in successive scans. The STD spectrum was then constructed by simple subtraction of both 2D ^1^H,^13^C-HSQC spectra. Blank 2D STD-^1^H,^13^C-HSQC experiment of the HA glycoproteins alone and glycans **3**-**5** were acquired as control.

^19^F CPMG NMR experiments were acquired on a 600 MHz spectrometer equipped with a Bruker selective ^19^F–^1^H decoupling (SEF) probe at 298 K, using samples containing the protein at the concentration of 10 μM, and the F-SLN **6** at 125 μM. Competitor glycans **7** and **2** were successively added to the sample. The standard CPMG Bruker pulse sequence was modified as described^30^. Twenty four (24) points were acquired with total echo times from 8 to 5200 ms, with τ = 2 ms. Data were analysed with the T1T2 relaxation module of Topspin3.5.

### In silico studies

#### MD simulations

When available, starting poses of the HAs were derived from X-ray crystal structures: HK68, A/HK/1/ 1968 H3N2 influenza virus hemagglutinin in complex with 6’-SLNLN (pdb code 6TZB)^24^. NL03, A/Wy/3/03 influenza virus hemagglutinin in complex with 6’-SLN (pdb code 6BKR)^26^. The starting poses of NL91, NL09, H3Sing and H3Sing (Y159A), were generated by superimposition of the model derived structure [http://3dflu.cent.uw.edu.pl/index.html]^31^ onto the X-ray crystal structure of the closest variant. The 3D coordinates of glycan **2** were generated by using the carbohydrate builder GLYCAM-web site [http://glycam.org]. The glycosidic torsion angles of the monosaccharides were maintained as observed by X-ray crystallography, while those not resolved were defined according to the lower energy values predicted by the GLYCAM-web modeling tool. The resulting poses were used as starting points for molecular dynamics (MD) simulations. The MD simulations were performed using the Amber16 program4 with the protein.ff14SB, the GLYCAM_06j-1 and the water.tip3p force fields parameters^32,33^. Next, the starting 3D geometries were placed into a 10 Å octahedral box of explicit TIP3P waters, and counter ions were added to maintain electro-neutrality. Two consecutive minimization steps were performed involving (1) only the water molecules and ions and (2) the whole system with a higher number of cycles, using the steepest descent algorithm. The system was subjected to two rapid molecular dynamic simulations (heating and equilibration). The equilibrated structures were the starting points for a final MD simulation at constant temperature (300 K) and pressure (1 atm). 1 μs molecular dynamic simulations without constraints were recorded, using an NPT ensemble with periodic boundary conditions, a cut-off of 10 Å, and the particle mesh Ewald method. A total of 500,000,000 molecular dynamics steps were run with a time step of 1 fs per step. Coordinates and energy values were recorded every 500000 steps (500 ps) for a total of 1000 MD models. The detailed analysis of the H-bond and CH-π interactions was performed along the MD trajectory using the cpptraj module included in Amber-Tools 16 package.

## Notes

The authors declare no competing financial interest.

## Acknowledgments

This research was supported by the Human Frontier Science Program Organization (HFSP) grant LT000747/2018-C (L.U.). R.P.d.V. is a recipient of an ERC starting grant from the European Commission (802780).

