## Supplementary material for "Probing Altered Receptor Specificities of Antigenically Drifting Human H3N2 Viruses by Chemoenzymatic Synthesis, NMR and Modeling": SI bioRxiv 5Apr2023 (from another version of this preprint: version 1, 2023-04-05)

**Table of Contents**

1. Compounds Preparation and Characterization
   1.1 NMR Spectra (Supplementary Figs. 1-3)
2. Synthesis of compound **6**
3. 2D STD-$^1$H, $^{13}$C-HSQC NMR spectra
   3.1 2D STD-$^1$H, $^{13}$C-HSQC NMR spectra of compound **4** with HK68, NL91 and NL03 (Supplementary Fig. 4).
   3.2 2D STD-$^1$H, $^{13}$C-HSQC NMR spectra of compound **5** with HK68, NL91 and NL03 (Supplementary Fig. 5).
4. Details of 2D STD NMR results for the interaction of compound **5** with HA NL03 protein (Supplementary Fig. 6).
5. Details of the intermolecular interactions between **2** and the HAs proteins as derived by molecular modelling (Supplementary Fig. 7).
6. Snapshoot of the **2**/HK68 MD trajectory (Supplementary Fig. 8).
7. Snapshoot of the **2**/NL91 MD trajectory (Supplementary Fig. 9).
8. Snapshoot of the **2**/NL03 MD trajectory (Supplementary Fig. 10).

### 1. Compound Preparation and Characterization

Compound **8**

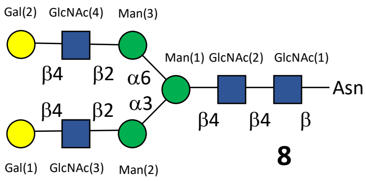


**8**

Compound **8** was prepared by a previously reported approach starting from Sialoglycopeptide [1].

$^1$H NMR (600 MHz, $D_2O$): δ = 5.03 (d, J < 2.0 Hz, 1H, H1-Man(2)), 4.99 (d, J = 9.8 Hz, 1H, H1-GlcNAc(1)), 4.83 (d, J < 2.0 Hz, 1H, H1-Man(3)), 4.69 (d, J = 2.6 Hz, 1H, H1-Man(1)), 4,53 (d, J = 7.6 Hz, 1H, H1-GlcNAc(2)), 4.50 (d, J = 7.8 Hz, 1H, H1-GlcNAc(3)), 4.47 (d, J = 7.8 Hz, 1H, H1-GlcNAc(4)), 4.38 (d, J = 7.7 Hz, 2H, H1-Gal(1), H1-Gal(2)), 4.17 (dd, J = 2.6 Hz, J = 6.5 Hz, 1H, H2-Man(1)), 4.11 (dd, J < 2.0 Hz, J = 2.6 Hz, H2-Man(2)), 4.00 (dd, J < 2.0 Hz, J = 2.6 Hz, H2-Man(3)), 3.97-3.30 (m, 52H, H2-H6-GlcNAc(1), H2-H6-GlcNAc(2), H3-H6-Man(1), H3-H6-Man(2), H3-H6-Man(3), H2-H6-GlcNAc(4), H2-H6-GlcNAc(3), H2-H6-Gal(1), H2-H6-Gal(2), αCH-Asn), 2.84-2.78 (m, 2H, βCH2-Asn), 1.99-1.95 (m, 12H, NHAc-GlcNAc(1), NHAc-GlcNAc(2), NHAc-GlcNAc(3), NHAc-GlcNAc(4) ppm.

$^{13}$C NMR (151 MHz, $D_2O$): d = 102.9, 101.3, 100.5, 99.5, 97.0, 80.4, 79.7, 78.7, 78.4, 78.1, 76.4, 76.2, 75.3, 75.2, 74.8, 74.4, 74.2, 73.5, 72.9, 72.3, 72.2, 72.0, 71.0, 70.1, 69.2, 68.6, 67.3, 65.9, 61.8, 61.5, 61.0, 60.0, 59.9, 55.0, 53.6, 50.9, 35.0, 22.4, 22.0 ppm.

ESI TOF-MS: for $C_{66}H_{110}N_6O_{48}$: m/z $[M-2H]^{2-}$ ; calcd: 876.3102; found: 876.3111.

Compound **9**

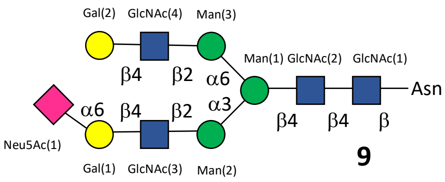


**9**

Compound **9** was prepared by a previously reported approach starting from compound **8** [2].

$^{1}$H NMR (600 MHz, $D_2O$): d = 5.03 (d, J < 2.0 Hz, 1H, H1-Man(2)), 4.99 (d, J = 9.8 Hz, 1H, H1-GlcNAc(1)), 4.83 (d, J < 2.0 Hz, 1H, H1-Man(3)), 4.69 (d, J = 2.6 Hz, 1H, H1-Man(1)), 4,53 (d, J = 7.6 Hz, 1H, H1-GlcNAc(2)), 4.50 (d, J = 7.8 Hz, 1H, H1-GlcNAc(3)), 4.47 (d, J = 7.8 Hz, 1H, H1-GlcNAc(4)), 4.38 (d, J = 7.7 Hz, 2H, H1-Gal(1), H1-Gal(2)), 4.17 (dd, J = 2.6 Hz, J = 6.5 Hz, 1H, H2-Man(1)), 4.11 (dd, J < 2.0 Hz, J = 2.6 Hz, H2-Man(2)), 4.00 (dd, J < 2.0 Hz, J = 2.6 Hz, H2-Man(3)), 3.97-3.30 (m, 59H, H2-H6-GlcNAc(1), H2-H6-GlcNAc(2), H3-H6-Man(1), H3-H6-Man(2), H3-H6-Man(3), H2-H6-GlcNAc(4), H2-H6-GlcNAc(3), H2-H6-Gal(1), H2-H6-Gal(2), H4-H9-Neu5Ac(1), αCH-Asn), 2.59 (dd, J = 12.2, J = 4.2, 1H, H3eq-Neu5Ac(1), 2.84-2.78 (m, 2H, βCH2-Asn), 1.99-1.95 (m, 15H, NHAc-GlcNAc(1), NHAc-GlcNAc(2), NHAc-GlcNAc(3), NHAc-GlcNAc(4), NHAc-Neu5Ac, 1.63 (t, J = 12.2, 1H, H3ax-Neu5Ac(1), ppm.

$^{13}$C NMR (151 MHz, $D_2O$): d = 102.9, 101.3, 100.5, 99.5, 97.0, 80.4, 79.7, 78.7, 78.4, 78.1, 76.4, 76.2, 75.3, 75.2, 74.8, 74.4, 74.2, 73.5, 72.9, 72.5, 72.3, 72.2, 72.0, 71.7, 71.0, 70.1, 69.2, 68.6, 68.3, 68.2, 67.3, 65.9, 63.3, 62.6, 61.8, 61.5, 61.0, 60.0, 59.9, 55.0, 53.6, 51.8, 50.9, 40.0, 35.0, 22.4, 22.0 ppm.

ESI TOF-MS: for $C_{77}H_{124}N_7O_{56}$: m/z $[M-2H]^{2-}$ ; calcd: 1021.8580; found: 1021.8575.

Compound **10**

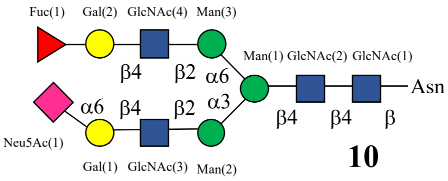


**10**

Compound **10** was prepared from **8** (12 mg, 6.8 µmol) using the general procedure for the installation of α2,6-Neu5Ac with ST6Gal1 and with Fut1 for the installation of α1,2 Fuc. After P2 column purification, **10** was obtained as a white solid (12.0 mg, 81%).

$^{1}$H NMR (600 MHz, $D_2O$): d = 5.22 (d, J = 2.0 Hz, 1H, H1-Fuc(1)), 5.03 (d, J < 2.0 Hz, 1H, H1-Man(2)), 4.99 (d, J = 9.8 Hz, 1H, H1-GlcNAc(1)), 4.83 (d, J < 2.0 Hz, 1H, H1-Man(3)), 4.69 (d, J = 2.6 Hz, 1H, H1-Man(1)), 4,53 (d, J = 7.6 Hz, 1H, H1-GlcNAc(2)), 4.50 (d, J = 7.8 Hz, 1H, H1-GlcNAc(3)), 4.47 (d, J = 7.8 Hz, 1H, H1-GlcNAc(4)), 4.38 (d, J = 7.7 Hz, 2H, H1-Gal(1), H1-Gal(2)), 4.17 (dd, J = 2.6 Hz, J = 6.5 Hz, 1H, H2-Man(1)), 4.13 (dd, J < 2.0 Hz, J = 6.7 Hz, 1H, H5-Fuc(1)), 4.11 (dd, J < 2.0 Hz, J = 2.6 Hz, H2-Man(2)), 4.00 (dd, J < 2.0 Hz, J = 2.6 Hz, H2-Man(3)), 3.97-3.30 (m, 62H, H2-H6-GlcNAc(1), H2-H6-GlcNAc(2), H3-H6-Man(1), H3-H6-Man(2), H3-H6-Man(3), H2-H6-GlcNAc(4), H2-H6-GlcNAc(3), H2-H6-Gal(1), H2-H6-Gal(2), H4-H9-Neu5Ac(1), H2-H4-Fuc(1), αCH-Asn), 2.59 (dd, J = 12.2, J = 4.2, 1H, H3eq-Neu5Ac(1), 2.84-2.78 (m, 2H, βCH2-Asn), 1.99-1.95 (m, 15H, NHAc-GlcNAc(1), NHAc-GlcNAc(2), NHAc-GlcNAc(3), NHAc-GlcNAc(4), NHAc-Neu5Ac, 1.63 (t, J = 12.2, 1H, H3ax-Neu5Ac(1), 1.15 (d, J = 6.7 Hz, 3H, H6-Fuc(1), ppm.

$^{13}$C NMR (151 MHz, $D_2O$): d = 102.9, 101.3, 100.5, 99.5, 99.4, 97.0, 80.4, 79.7, 78.7, 78.4, 78.1, 76.5, 76.4, 76.2, 75.3, 75.2, 74.8, 74.4, 74.2, 73.5, 72.9, 72.5, 72.3, 72.2, 72.0, 71.7, 71.0, 70.1, 69.5, 69.3, 69.2, 68.6, 68.3, 68.2, 67.3, 66.9, 65.9, 63.3, 62.6, 61.8, 61.5, 61.0, 60.0, 59.9, 55.0, 53.6, 51.8, 50.9, 40.0, 35.0, 22.4, 22.0, 15.5, ppm.

ESI TOF-MS: for $C_{83}H_{135}N_7O_{60}$: m/z $[M-2H]^{2-}$ ; calcd: 1095.3908; found: 1095.3925.

**Preparation of compound 11**

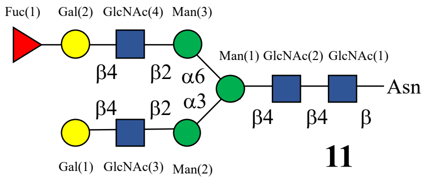


**11**

Compound **11** was prepared from **10** (12 mg, 5.4 µmol) by acid mediated hydrolysis of the α2,6-Neu5Ac according to the general procedure. Compound **10** was dissolved in 4 ml aqueous solution of acetic acid (2 M) and kept at 65 °C for 24 h. The solvent was removed by an $N_2$ flow. After purification by HPLC over an HILIC column, **11** was obtained as a white solid (6.5 mg, 45%).

$^{1}$H NMR (600 MHz, $D_2O$): d = 5.22 (d, J = 2.0 Hz, 1H, H1-Fuc(1)), 5.03 (d, J < 2.0 Hz, 1H, H1-Man(2)), 4.99 (d, J = 9.8 Hz, 1H, H1-GlcNAc(1)), 4.83 (d, J < 2.0 Hz, 1H, H1-Man(3)), 4.69 (d, J = 2.6 Hz, 1H, H1-Man(1)), 4,53 (d, J = 7.6 Hz, 1H, H1-GlcNAc(2)), 4.50 (d, J = 7.8 Hz, 1H, H1-GlcNAc(3)), 4.47 (d, J = 7.8 Hz, 1H, H1-GlcNAc(4)), 4.46 (d, J = 7.7 Hz, 1H, H1-Gal(2)), 4.38 (d, J = 7.6 Hz, 1H, H1-Gal(1)), 4.17 (dd, J = 2.6 Hz, J = 6.5 Hz, 1H, H2-Man(1)), 4.13 (dd, J < 2.0 Hz, J = 6.7 Hz, 1H, H5-Fuc(1)), 4.11 (dd, J < 2.0 Hz, J = 2.6 Hz, H2-Man(2)), 4.00 (dd, J < 2.0 Hz, J = 2.6 Hz, H2-Man(3)), 3.97-3.30 (m, 55H, H2-H6-GlcNAc(1), H2-H6-GlcNAc(2), H3-H6-Man(1), H3-H6-Man(2), H3-H6-Man(3), H2-H6-GlcNAc(4), H2-H6-GlcNAc(3), H2-H6-Gal(1), H2-H6-Gal(2), H2-H4-Fuc(1), αCH-Asn), 2.84-2.78 (m, 2H, βCH2-

Asn), 1.99-1.95 (m, 12H, NHAc-GlcNAc(1), NHAc-GlcNAc(2), NHAc-GlcNAc(3), NHAc-GlcNAc(4), 1.15 (d, J = 6.7 Hz, 3H, H6-Fuc(1), ppm.

$^{13}$C NMR (151 MHz, $D_2O$): d = 102.9, 101.3, 100.5, 99.5, 99.4, 97.0, 80.4, 79.7, 78.7, 78.4, 78.1, 76.5, 76.4, 76.2, 75.3, 75.2, 74.8, 74.4, 74.2, 73.5, 72.9, 72.3, 72.2, 72.0, 71.0, 70.1, 69.5, 69.3, 69.2, 68.6, 68.2, 67.3, 66.9, 65.9, 62.6, 61.8, 61.5, 61.0, 60.0, 59.9, 55.0, 53.6, 50.9, 35.0, 22.4, 22.0, 15.5, ppm.

ESI TOF-MS: for $C_{72}H_{118}N_6O_{52}$: m/z $[M-2H]^{2-}$ ; calcd: 949.3392; found: 949.3401.

### Preparation of compounds 12 and 17

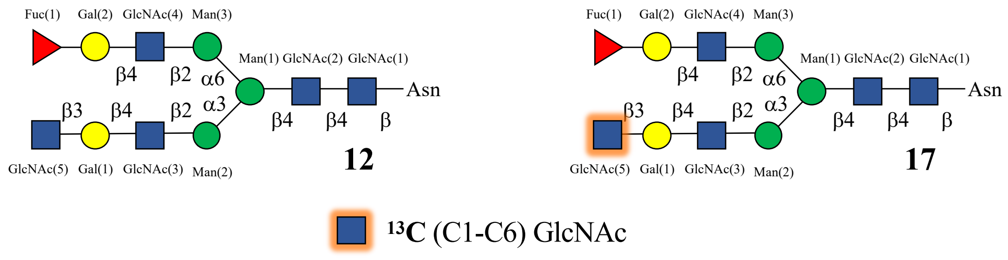


Compounds **12** and **17** were prepared by a previously reported approach starting from compound **11**.[2]

$^1$H NMR (600 MHz, $D_2O$): d = 5.22 (d, J = 2.0 Hz, 1H, H1-Fuc(1)), 5.03 (d, J < 2.0 Hz, 1H, H1-Man(2)), 4.99 (d, J = 9.8 Hz, 1H, H1-GlcNAc(1)), 4.83 (d, J < 2.0 Hz, 1H, H1-Man(3)), 4.69 (d, J = 2.6 Hz, 1H, H1-Man(1)), 4.59 (d, J = 9.6 Hz, 1H, H1-GlcNAc(5)), 4,53 (d, J = 7.6 Hz, 1H, H1-GlcNAc(2)), 4.50 (d, J = 7.8 Hz, 1H, H1-GlcNAc(3)), 4.47 (d, J = 7.8 Hz, 1H, H1-GlcNAc(4)), 4.46 (d, J = 7.7 Hz, 1H, H1-Gal(2)), 4.38 (d, J = 7.6 Hz, 1H, H1-Gal(1)), 4.17 (dd, J = 2.6 Hz, J = 6.5 Hz, 1H, H2-Man(1)), 4.13 (dd, J < 2.0 Hz, J = 6.7 Hz, 1H, H5-Fuc(1)), 4.11 (dd, J < 2.0 Hz, J = 2.6 Hz, H2-Man(2)), 4.00 (dd, J < 2.0 Hz, J = 2.6 Hz, H2-Man(3)), 3.97-3.30 (m, 61H, H2-H6-GlcNAc(1), H2-H6-GlcNAc(2), H3-H6-Man(1), H3-H6-Man(2), H3-H6-Man(3), H2-H6-GlcNAc(4), H2-H6-GlcNAc(3), H2-H6-Gal(1), H2-H6-Gal(2), H2-H4-Fuc(1), H2-H6-GlcNAc(5), αCH-Asn), 2.84-2.78 (m, 2H, βCH2-Asn), 1.99-1.95 (m, 15H, NHAc-GlcNAc(1), NHAc-GlcNAc(2), NHAc-GlcNAc(3), NHAc-GlcNAc(4), NHAc-GlcNAc(5) 1.15 (d, J = 6.7 Hz, 3H, H6-Fuc(1), ppm.

$^{13}$C NMR (151 MHz, $D_2O$): d = 102.9, 102.8, 101.3, 100.5, 100.2, 99.5, 99.4, 97.0, 82.0, 80.4, 79.7, 78.7, 78.4, 78.1, 76.5, 76.4, 76.2, 75.3, 75.2, 74.8, 74.4, 74.2, 73.5, 72.9, 72.3, 72.2, 72.0, 71.0, 70.1, 70.0, 69.5, 69.4, 69.3, 69.2, 68.6, 68.3, 68.2, 67.3, 66.9, 65.9, 62.6, 61.8, 61.5, 61.0, 60.0, 59.9, 55.7, 55.0, 53.6, 50.9, 35.0, 22.4, 22.0, 15.5, ppm.

ESI TOF-MS: for $C_{80}H_{133}N_7O_{57}$: m/z $[M-2H]^{2-}$ ; (**12**) calcd: 1050.8789; found: 1050.8767. (**17**) calcd: 1053.8890; found: 1053.8868.

### Preparation of compounds 13 and 18

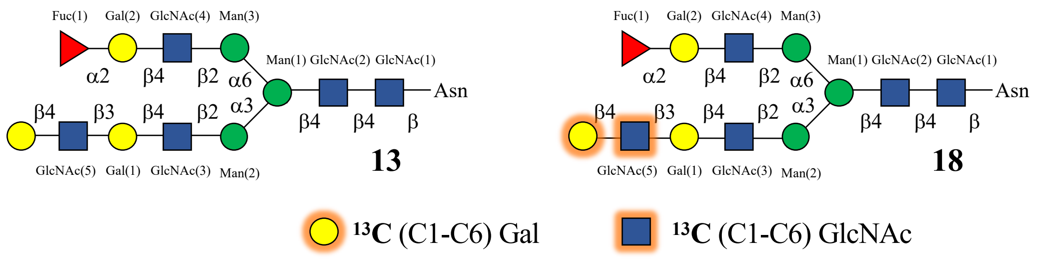


Compound **13** was prepared from **11** (5.0 mg, 2.6 µmol) using the general procedure for the installation of β1,3-GlcNAc with B3GnT2 and with B4GalT1 for the installation of β1,4-Gal. After P2 column purification, **13** was obtained as a white solid (1.9 mg, 82%).

Compound **18** was prepared from **11** (1.5 mg, 0.8 µmol) using the general procedure for the installation of β1,3-GlcNAc with B3GnT2 and B4GalT1 for the installation of β1,4-Gal. After P2 column purification, **18** was obtained as a white solid (1.5 mg, 83%).

$^{1}$H NMR (600 MHz, $D_2O$): d = 5.22 (d, J = 2.0 Hz, 1H, H1-Fuc(1)), 5.03 (d, J < 2.0 Hz, 1H, H1-Man(2)), 4.99 (d, J = 9.8 Hz, 1H, H1-GlcNAc(1)), 4.83 (d, J < 2.0 Hz, 1H, H1-Man(3)), 4.69 (d, J = 2.6 Hz, 1H, H1-Man(1)), 4.59 (d, J = 9.6 Hz, 1H, H1-GlcNAc(5)), 4,53 (d, J = 7.6 Hz, 1H, H1-GlcNAc(2)), 4.50 (d, J = 7.8 Hz, 1H, H1-GlcNAc(3)), 4.47 (d, J = 7.8 Hz, 1H, H1-GlcNAc(4)), 4.46 (d, J = 7.7 Hz, 1H, H1-Gal(2)), 4.38 (d, J = 7.6 Hz, 2H, H1-Gal(1), H1-Gal(3)), 4.17 (dd, J = 2.6 Hz, J = 6.5 Hz, 1H, H2-Man(1)), 4.13 (dd, J < 2.0 Hz, J = 6.7 Hz, 1H, H5-Fuc(1)), 4.11 (dd, J < 2.0 Hz, J = 2.6 Hz, H2-Man(2)), 4.00 (dd, J < 2.0 Hz, J = 2.6 Hz, H2-Man(3)), 3.97-3.30 (m, 67H, H2-H6-GlcNAc(1), H2-H6-GlcNAc(2), H3-H6-Man(1), H3-H6-Man(2), H3-H6-Man(3), H2-H6-GlcNAc(4), H2-H6-GlcNAc(3), H2-H6-Gal(1), H2-H6-Gal(2), H2-H4-Fuc(1), H2-H6-GlcNAc(5), H2-H6-Gal(3), αCH-Asn), 2.84-2.78 (m, 2H, βCH2-Asn), 1.99-1.95 (m, 15H, NHAc-GlcNAc(1), NHAc-GlcNAc(2), NHAc-GlcNAc(3), NHAc-GlcNAc(4), NHAc-GlcNAc(5) 1.15 (d, J = 6.7 Hz, 3H, H6-Fuc(1), ppm.

$^{13}$C NMR (151 MHz, $D_2O$): d = 102.9, 102.8, 101.3, 100.5, 100.2, 99.5, 99.4, 97.0, 82.0, 80.4, 79.7, 78.7, 78.4, 78.1, 77.8, 76.5, 76.4, 76.2, 75.3, 75.2, 74.8, 74.4, 74.3, 74.2, 73.5, 72.9, 72.3, 72.2, 72.0, 71.0, 70.1, 70.0, 69.5, 69.4, 69.3, 69.2, 68.6, 68.3, 68.2, 67.3, 66.9, 65.9, 62.6, 61.8, 61.5, 61.0, 60.0, 59.9, 59.7, 55.7, 55.0, 53.6, 50.9, 35.0, 22.4, 22.0, 15.5, ppm.

ESI TOF-MS: for $C_{86}H_{142}N_7O_{62}$: m/z $[M-2H]^{2-}$ ; (**13**) calcd: 1131.4014; found: 1131.4002. (**18**) calcd: 1137.4215; found: 1137.4208.

### Preparation of compounds 14 and 19

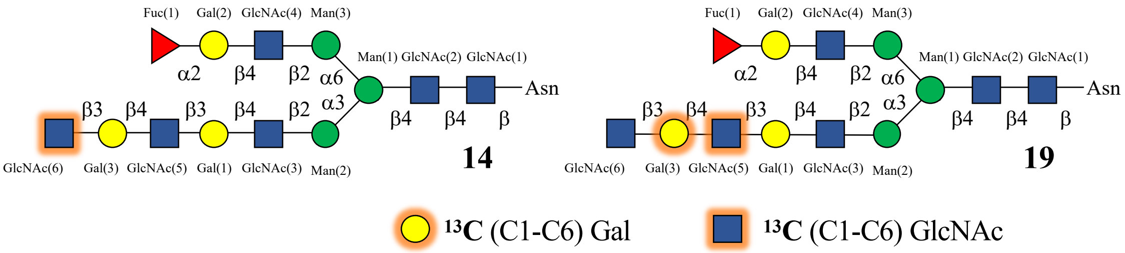


Compounds **14** and **19** were prepared by a previously reported approach starting from compound **13** and **18**, respectively.[2]

$^{1}$H NMR (600 MHz, $D_2O$): d = 5.22 (d, J = 2.0 Hz, 1H, H1-Fuc(1)), 5.03 (d, J < 2.0 Hz, 1H, H1-Man(2)), 4.99 (d, J = 9.8 Hz, 1H, H1-GlcNAc(1)), 4.83 (d, J < 2.0 Hz, 1H, H1-Man(3)), 4.69 (d, J = 2.6 Hz, 1H, H1-Man(1)), 4.59 (d, J = 9.6 Hz, 2H, H1-GlcNAc(5), H1-GlcNAc(6)), 4,53 (d, J = 7.6 Hz, 1H, H1-GlcNAc(2)), 4.50 (d, J = 7.8 Hz, 1H, H1-GlcNAc(3)), 4.47 (d, J = 7.8 Hz, 1H, H1-GlcNAc(4)), 4.46 (d, J = 7.7 Hz, 1H, H1-Gal(2)), 4.38 (d, J = 7.6 Hz, 2H, H1-Gal(1), H1-Gal(3)), 4.17 (dd, J = 2.6 Hz, J = 6.5 Hz, 1H, H2-Man(1)), 4.13 (dd, J < 2.0 Hz, J = 6.7 Hz, 1H, H5-Fuc(1)), 4.11 (dd, J < 2.0 Hz, J = 2.6 Hz, H2-Man(2)), 4.06 (dd, J < 2.0 Hz, J = 2.4 Hz, 2H, H4-Gal(1), H4-Gal(3)), 4.00 (dd, J < 2.0 Hz, J = 2.6 Hz, H2-Man(3)), 3.97-3.30 (m, 72H, H2-H6-GlcNAc(1), H2-H6-GlcNAc(2), H3-H6-Man(1), H3-H6-Man(2), H3-H6-Man(3), H2-H6-GlcNAc(4), H2-H6-GlcNAc(3), H2-H6-Gal(1), H2-H6-Gal(2), H2-H4-Fuc(1), H2-H6-GlcNAc(5), H3-H6-Gal(3), H2-H6-GlcNAc(6), αCH-Asn), 2.84-2.78 (m, 2H, βCH2-Asn), 1.99-1.95 (m, 18H, NHAc-GlcNAc(1), NHAc-GlcNAc(2), NHAc-GlcNAc(3), NHAc-GlcNAc(4), NHAc-GlcNAc(5), NHAc-GlcNAc(6), 1.15 (d, J = 6.7 Hz, 3H, H6-Fuc(1), ppm.

$^{13}$C NMR (151 MHz, $D_2O$): d = 102.9, 102.8, 101.3, 100.5, 100.2, 99.5, 99.4, 97.0, 82.0, 80.4, 79.7, 78.7, 78.4, 78.1, 77.8, 76.5, 76.4, 76.2, 75.3, 75.2, 74.8, 74.4, 74.3, 74.2, 73.5, 72.9, 72.3, 72.2, 72.0, 71.0, 70.1, 70.0, 69.5, 69.4, 69.3, 69.2, 68.6, 68.3, 68.2, 67.3, 66.9, 65.9, 62.6, 61.8, 61.5, 61.0, 60.0, 59.9, 59.7, 55.7, 55.0, 53.6, 50.9, 35.0, 22.4, 22.0, 15.5, ppm.

ESI TOF-MS: for $C_{94}H_{156}N_8O_{67}$: m/z $[M-2H]^{2-}$ ; (**14**) calcd: 1235.9511; found: 1235.9492. (**19**) calcd: 1238.9612; found: 1238.9589.

### Preparation of compounds 15 and 20

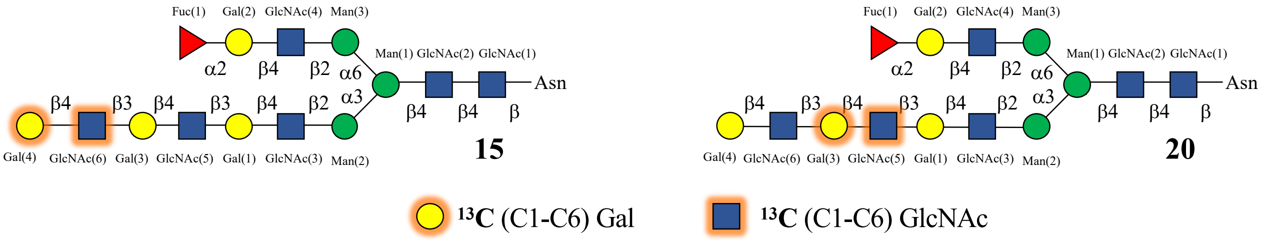


Compounds **15** and **20** were prepared by a previously reported approach starting from compound **14** and **19**, respectively.[2]

$^{1}$H NMR (600 MHz, $D_2O$): d = 5.22 (d, J = 2.0 Hz, 1H, H1-Fuc(1)), 5.03 (d, J < 2.0 Hz, 1H, H1-Man(2)), 4.99 (d, J = 9.8 Hz, 1H, H1-GlcNAc(1)), 4.83 (d, J < 2.0 Hz, 1H, H1-Man(3)), 4.69 (d, J = 2.6 Hz, 1H, H1-Man(1)), 4.59 (d, J = 9.6 Hz, 2H, H1-GlcNAc(5), H1-GlcNAc(6)), 4,53 (d, J = 7.6 Hz, 1H, H1-GlcNAc(2)),

4.50 (d, J = 7.8 Hz, 1H, H1-GlcNAc(3)), 4.47 (d, J = 7.8 Hz, 1H, H1-GlcNAc(4)), 4.46 (d, J = 7.7 Hz, 1H, H1-Gal(2)), 4.38 (d, J = 7.6 Hz, 3H, H1-Gal(1), H1-Gal(3), H1-Gal(4)), 4.17 (dd, J = 2.6 Hz, J = 6.5 Hz, 1H, H2-Man(1)), 4.13 (dd, J < 2.0 Hz, J = 6.7 Hz, 1H, H5-Fuc(1)), 4.11 (dd, J < 2.0 Hz, J = 2.6 Hz, H2-Man(2)), 4.06 (dd, J < 2.0 Hz, J = 2.4 Hz, 2H, H4-Gal(1), H4-Gal(3)), 4.00 (dd, J < 2.0 Hz, J = 2.6 Hz, H2-Man(3)), 3.97-3.30 (m, 78H, H2-H6-GlcNAc(1), H2-H6-GlcNAc(2), H3-H6-Man(1), H3-H6-Man(2), H3-H6-Man(3), H2-H6-GlcNAc(4), H2-H6-GlcNAc(3), H2-H6-Gal(1), H2-H6-Gal(2), H2-H4-Fuc(1), H2-H6-GlcNAc(5), H3-H6-Gal(3), H2-H6-GlcNAc(6), H2-H6-Gal(4), αCH-Asn), 2.84-2.78 (m, 2H, βCH2-Asn), 1.99-1.95 (m, 18H, NHAc-GlcNAc(1), NHAc-GlcNAc(2), NHAc-GlcNAc(3), NHAc-GlcNAc(4), NHAc-GlcNAc(5), NHAc-GlcNAc(6), 1.15 (d, J = 6.7 Hz, 3H, H6-Fuc(1), ppm.

$^{13}$C NMR (151 MHz, $D_2O$): d = 102.9, 102.8, 101.3, 100.5, 100.2, 99.5, 99.4, 97.0, 82.0, 80.4, 79.7, 78.7, 78.4, 78.1, 77.8, 76.5, 76.4, 76.2, 75.3, 75.2, 74.8, 74.4, 74.3, 74.2, 73.5, 72.9, 72.3, 72.2, 72.0, 71.0, 70.1, 70.0, 69.5, 69.4, 69.3, 69.2, 68.6, 68.3, 68.2, 67.3, 66.9, 65.9, 62.6, 61.8, 61.5, 61.0, 60.0, 59.9, 59.7, 55.7, 55.0, 53.6, 50.9, 35.0, 22.4, 22.0, 15.5, ppm.

ESI TOF-MS: for $C_{100}H_{166}N_8O_{72}$: m/z $[M-2H]^{2-}$ ; (**15**) calcd: 1320.4915; found: 1320.4887. (**20**) calcd: 1320.4915; found: 1320.4902.

### Preparation of compounds 16 and 21

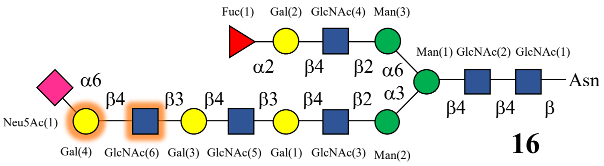


**16**

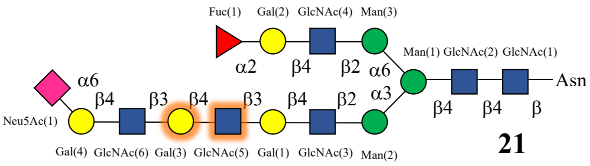


**21**

$^{13}$C (C1-C6) Gal    $^{13}$C (C1-C6) GlcNAc

Compound **16** was prepared from **13** (1.9 mg, 0.84 µmol) using the general procedure for the installation of β1,3-GlcNAc with B3GnT2, and B4GalT1 for the installation of β1,4-Gal and ST6Gal1 for the installation of the α2,6-Neu5Ac. After P2 column purification, **16** was obtained as a white solid (0.7 mg, 29% over three steps).

Compound **21** was prepared from **18** (1.5 mg, 0.66 µmol) using the general procedure for the installation of β1,3-GlcNAc with B3GnT2 and B4GalT1 for the installation of β1,4-Gal and ST6Gal1 for the installation of the α2,6-Neu5Ac. After P2 column purification, **21** was obtained as a white solid (0.55 mg, 30% over three steps.

$^{1}$H NMR (600 MHz, $D_2O$): d = 5.22 (d, J = 2.0 Hz, 1H, H1-Fuc(1)), 5.03 (d, J < 2.0 Hz, 1H, H1-Man(2)), 4.99 (d, J = 9.8 Hz, 1H, H1-GlcNAc(1)), 4.83 (d, J < 2.0 Hz, 1H, H1-Man(3)), 4.69 (d, J = 2.6 Hz, 1H, H1-Man(1)), 4.59 (d, J = 9.6 Hz, 2H, H1-GlcNAc(5), H1-GlcNAc(6)), 4,53 (d, J = 7.6 Hz, 1H, H1-GlcNAc(2)), 4.50 (d, J = 7.8 Hz, 1H, H1-GlcNAc(3)), 4.47 (d, J = 7.8 Hz, 1H, H1-GlcNAc(4)), 4.46 (d, J = 7.7 Hz, 1H, H1-Gal(2)), 4.38 (d, J = 7.6 Hz, 3H, H1-Gal(1), H1-Gal(3)), 4.35 (d, J = 7.8 Hz, H1-Gal(4), 4.17 (dd, J = 2.6 Hz, J = 6.5 Hz, 1H, H2-Man(1)), 4.13 (dd, J < 2.0 Hz, J = 6.7 Hz, 1H, H5-Fuc(1)), 4.11 (dd, J < 2.0 Hz, J = 2.6 Hz, H2-Man(2)), 4.06 (dd, J < 2.0 Hz, J = 2.4 Hz, 3H, H4-Gal(1), H4-Gal(3), H4-Gal(4)), 4.00 (dd, J < 2.0 Hz, J = 2.6 Hz, H2-Man(3)), 3.97-3.30 (m, 85H, H2-H6-GlcNAc(1), H2-H6-GlcNAc(2), H3-H6-Man(1), H3-H6-Man(2), H3-H6-Man(3), H2-H6-GlcNAc(4), H2-H6-GlcNAc(3), H2-H6-Gal(1), H2-H6-

Gal(2), H2-H4-Fuc(1), H2-H6-GlcNAc(5), H3-H6-Gal(3), H2-H6-GlcNAc(6), H2-H6-Gal(4), H4-H9-Neu5Ac(1), αCH-Asn), 2.84-2.78 (m, 2H, βCH2-Asn), 2.59 (dd, J = 12.2, J = 4.2, 1H, H3eq-Neu5Ac(1), 1.99-1.95 (m, 21H, NHAc-GlcNAc(1), NHAc-GlcNAc(2), NHAc-GlcNAc(3), NHAc-GlcNAc(4), NHAc-GlcNAc(5), NHAc-GlcNAc(6), NHAc-Neu5Ac(1), NHAc-Neu5Ac, 1.63 (t, J = 12.2, 1H, H3ax-Neu5Ac(1), 1.15 (d, J = 6.7 Hz, 3H, H6-Fuc(1), ppm.

$^{13}$C NMR (151 MHz, $D_2O$): d = 102.9, 102.8, 101.3, 100.5, 100.2, 99.5, 99.4, 97.0, 82.0, 80.4, 79.7, 78.7, 78.4, 78.1, 77.8, 76.5, 76.4, 76.2, 75.3, 75.2, 74.8, 74.4, 74.3, 74.2, 73.5, 72.9, 72.3, 72.2, 72.0, 71.0, 70.1, 70.0, 69.5, 69.4, 69.3, 69.2, 68.6, 68.3, 68.2, 67.3, 66.9, 65.9, 62.6, 61.8, 61.5, 61.0, 60.0, 59.9, 59.7, 55.7, 55.0, 53.6, 50.9, 35.0, 22.4, 22.0, 15.5, ppm.

ESI TOF-MS: for $C_{111}H_{182}N_9O_{80}$: m/z $[M-2H]^{2-}$ ; (**16**) calcd: 1466.0370; found: 1465.9885. (**21**) calcd: 1466.0370; found: 1466.0366.

### Preparation of compound 22

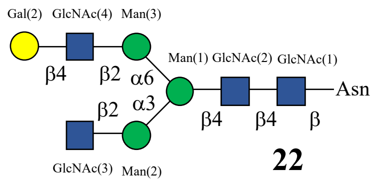


**22**

Compound **22** was prepared from **8** (5.0 mg, 2.8 μmol) by a previously reported procedure for the selective cleavage of galactoside with *E. coli* β-galactosidase.[2] After HPLC purification using a HILIC column, compound **22** was obtained as a white solid (3.1 mg, 71%).

$^{1}$H NMR (600 MHz, $D_2O$): d = 5.03 (d, J < 2.0 Hz, 1H, H1-Man(2)), 4.99 (d, J = 9.8 Hz, 1H, H1-GlcNAc(1)), 4.83 (d, J < 2.0 Hz, 1H, H1-Man(3)), 4.69 (d, J = 2.6 Hz, 1H, H1-Man(1)), 4,53 (d, J = 7.6 Hz, 1H, H1-GlcNAc(2)), 4.50 (d, J = 7.8 Hz, 1H, H1-GlcNAc(3)), 4.47 (d, J = 7.8 Hz, 1H, H1-GlcNAc(4)), 4.38 (d, J = 7.7 Hz, 1H, H1-Gal(2)), 4.17 (dd, J = 2.6 Hz, J = 6.5 Hz, 1H, H2-Man(1)), 4.11 (dd, J < 2.0 Hz, J = 2.6 Hz, H2-Man(2)), 4.00 (dd, J < 2.0 Hz, J = 2.6 Hz, H2-Man(3)), 3.97-3.30 (m, 46H, H2-H6-GlcNAc(1), H2-H6-GlcNAc(2), H3-H6-Man(1), H3-H6-Man(2), H3-H6-Man(3), H2-H6-GlcNAc(4), H2-H6-GlcNAc(3), H2-H6-Gal(2), αCH-Asn), 2.84-2.78 (m, 2H, βCH2-Asn), 1.99-1.95 (m, 12H, NHAc-GlcNAc(1), NHAc-GlcNAc(2), NHAc-GlcNAc(3), NHAc-GlcNAc(4) ppm.

$^{13}$C NMR (151 MHz, $D_2O$): d = 102.9, 101.3, 100.5, 99.5, 97.0, 80.4, 79.7, 78.7, 78.4, 78.1, 76.4, 76.2, 75.3, 75.2, 74.4, 74.2, 73.5, 72.9, 72.3, 72.2, 72.0, 71.0, 70.1, 69.2, 68.6, 67.3, 65.9, 61.8, 61.5, 61.0, 60.0, 55.0, 53.6, 50.9, 35.0, 22.4, 22.0 ppm.

ESI TOF-MS: for $C_{60}H_{100}N_6O_{43}$: m/z $[M-2H]^{2-}$ ; calcd: 795.2838; found: 795.2825.

### Preparation of compound 23

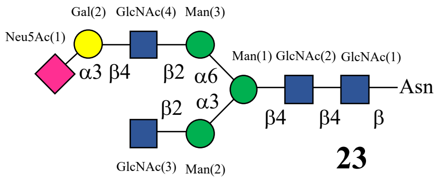


**23**

Compound **23** was prepared from **22** (3.0 mg, 1.9 μmol) using the general procedure for the installation of α2,3-Neu using ST3Gal4. After P2 column purification, compound **23** was obtained as a white solid (3.4 mg, 95%).

$^{1}$H NMR (600 MHz, $D_2O$): d = 5.03 (d, J < 2.0 Hz, 1H, H1-Man(2)), 4.99 (d, J = 9.8 Hz, 1H, H1-GlcNAc(1)), 4.83 (d, J < 2.0 Hz, 1H, H1-Man(3)), 4.69 (d, J = 2.6 Hz, 1H, H1-Man(1)), 4,53 (d, J = 7.6 Hz, 1H, H1-GlcNAc(2)), 4.50 (d, J = 7.8 Hz, 1H, H1-GlcNAc(3)), 4.47 (d, J = 7.8 Hz, 1H, H1-GlcNAc(4)), 4.43 (d, J = 7.7 Hz, 1H, H1-Gal(2)), 4.17 (dd, J = 2.6 Hz, J = 6.5 Hz, 1H, H2-Man(1)), 4.11 (dd, J < 2.0 Hz, J = 2.6 Hz, H2-Man(2)), 4.00 (dd, J < 2.0 Hz, J = 2.6 Hz, H2-Man(3)), 3.99 (dd, J < 2.0 Hz, J = 2.5 Hz, H4-Gal(2)), 3.97-3.30 (m, 53H, H2-H6-GlcNAc(1), H2-H6-GlcNAc(2), H3-H6-Man(1), H3-H6-Man(2), H3-H6-Man(3), H2-H6-GlcNAc(4), H2-H6-GlcNAc(3), H2-H3, H5-H6-Gal(2), H4-H9 Neu5Ac, αCH-Asn), 2.84-2.78 (m, 2H, βCH2-Asn), 2.64 (dd, J = 12.4, J = 4.6, H3eq-Neu5Ac), 1.99-1.95 (m, 15H, NHAc-GlcNAc(1), NHAc-GlcNAc(2), NHAc-GlcNAc(3), NHAc-GlcNAc(4), NHAc-Neu5Ac, 1.68 (t, J = 12.4, H3ax-Neu5Ac) ppm.

$^{13}$C NMR (151 MHz, $D_2O$): d = 102.6, 101.3, 100.5, 99.5, 97.0, 80.4, 79.7, 78.7, 78.4, 78.1, 76.4, 76.2, 75.4, 75.2, 74.4, 74.2, 73.5, 72.8, 72.3, 72.2, 72.0, 71.7, 71.0, 70.1, 69.3, 68.6, 68.3, 67.6, 65.9, 62.3, 61.8, 61.5, 61.0, 60.0, 55.0, 53.6, 51.7, 50.9, 39.6, 35.0, 22.4, 22.0 ppm.

ESI TOF-MS: for $C_{71}H_{117}N_7O_{51}$: m/z $[M-2H]^{2-}$ ; calcd: 940.8315; found: 940.8298.

### Preparation of compound 24

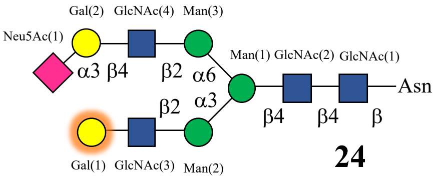


**24**

$^{13}$C (C1-C6) Gal

Compound **24** was prepared from **23** (3.4 mg, 1.8 µmol) using the general procedure for the installation of β1,3-Gal using B4GalT1. After P2 column purification, compound **24** was obtained as a white solid (2.9 mg, 81%).

$^1$H NMR (600 MHz, $D_2O$): d = 5.03 (d, J < 2.0 Hz, 1H, H1-Man(2)), 4.99 (d, J = 9.8 Hz, 1H, H1-GlcNAc(1)), 4.83 (d, J < 2.0 Hz, 1H, H1-Man(3)), 4.69 (d, J = 2.6 Hz, 1H, H1-Man(1)), 4,53 (d, J = 7.6 Hz, 1H, H1-GlcNAc(2)), 4.47 (d, J = 7.8 Hz, 2H, H1-GlcNAc(3), H1-GlcNAc(4)), 4.43 (d, J = 7.7 Hz, 1H, H1-Gal(2)), 4.38 (d, J = 7.6 Hz, 1H, H1-Gal(1)), 4.17 (dd, J = 2.6 Hz, J = 6.5 Hz, 1H, H2-Man(1)), 4.11 (dd, J < 2.0 Hz, J = 2.6 Hz, H2-Man(2)), 4.00 (dd, J < 2.0 Hz, J = 2.6 Hz, H2-Man(3)), 3.99 (dd, J < 2.0 Hz, J = 2.5 Hz, H4-Gal(2)), 3.97-3.30 (m, 59H, H2-H6-GlcNAc(1), H2-H6-GlcNAc(2), H3-H6-Man(1), H3-H6-Man(2), H3-H6-Man(3), H2-H6-GlcNAc(4), H2-H6-GlcNAc(3), H2-H3, H5-H6-Gal(2), H2-H6 -Gal(1), H4-H9 Neu5Ac, αCH-Asn), 2.84-2.78 (m, 2H, βCH2-Asn), 2.64 (dd, J = 12.4, J = 4.6, H3eq-Neu5Ac), 1.99-1.95 (m, 15H, NHAc-GlcNAc(1), NHAc-GlcNAc(2), NHAc-GlcNAc(3), NHAc-GlcNAc(4), NHAc-Neu5Ac, 1.68 (t, J = 12.4, H3ax-Neu5Ac) ppm.

$^{13}$C NMR (151 MHz, $D_2O$): d = 102.9, 102.6, 101.3, 100.5, 99.5, 97.0, 80.4, 79.7, 78.7, 78.4, 78.1, 76.4, 76.2, 75.4, 75.3, 75.2, 74.4, 74.2, 73.5, 72.8, 72.3, 72.2, 72.0, 71.7, 71.0, 70.1, 69.3, 68.6, 68.3, 67.6, 65.9, 62.3, 61.8, 61.5, 61.0, 60.0, 55.0, 53.6, 51.7, 50.9, 39.6, 35.0, 22.4, 22.0 ppm.

ESI TOF-MS: for $C_{77}H_{127}N_7O_{56}$: m/z $[M-2H]^{2-}$ ; calcd: 1024.8680; found: 1024.8655.

### Preparation of compound 25

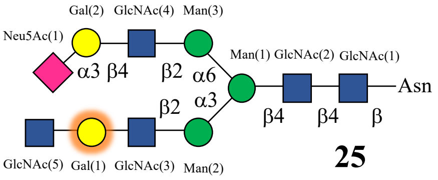


**25**

$^{13}$C (C1-C6) Gal

Compound **25** was prepared from **24** (2.9 mg, 1.45 µmol) using the general procedure for the installation of β1,4-GlcNAc using B3GnT2. After P2 column purification, compound **25** was obtained as a white solid (2.6 mg, 1.2 µmol, 83%).

$^1$H NMR (600 MHz, $D_2O$): d = 5.03 (d, J < 2.0 Hz, 1H, H1-Man(2)), 4.99 (d, J = 9.8 Hz, 1H, H1-GlcNAc(1)), 4.83 (d, J < 2.0 Hz, 1H, H1-Man(3)), 4.69 (d, J = 2.6 Hz, 1H, H1-Man(1)), 4.59 (d, J = 9.7 Hz, 1H, H1-GlcNAc(5)), 4,53 (d, J = 7.6 Hz, 1H, H1-GlcNAc(2)), 4.47 (d, J = 7.8 Hz, 2H, H1-GlcNAc(3), H1-GlcNAc(4)), 4.43 (d, J = 7.7 Hz, 1H, H1-Gal(2)), 4.38 (d, J = 7.6 Hz, 1H, H1-Gal(1)), 4.17 (dd, J = 2.6 Hz, J = 6.5 Hz, 1H, H2-Man(1)), 4.11 (dd, J < 2.0 Hz, J = 2.6 Hz, H2-Man(2)), 4.06 (dd, J < 2.0 Hz, J = 2.6 Hz, H4-Gal(1)), 4.00 (dd, J < 2.0 Hz, J = 2.6 Hz, H2-Man(3)), 3.99 (dd, J < 2.0 Hz, J = 2.5 Hz, H4-Gal(2)), 3.97-3.30 (m, 65H, H2-H6-GlcNAc(1), H2-H6-GlcNAc(2), H3-H6-Man(1), H3-H6-Man(2), H3-H6-Man(3), H2-H6-GlcNAc(4), H2-H6-GlcNAc(3), H2-H3, H5-H6-Gal(2), H2-H6 -Gal(1), H2-H6-GlcNAc(5), H4-H9 Neu5Ac, αCH-Asn), 2.84-2.78 (m, 2H, βCH2-Asn), 2.64 (dd, J = 12.4, J = 4.6, H3eq-Neu5Ac), 1.99-1.95 (m, 18H, NHAc-GlcNAc(1), NHAc-GlcNAc(2), NHAc-GlcNAc(3), NHAc-GlcNAc(4), NHAc-GlcNAc(5), NHAc-Neu5Ac, 1.68 (t, J = 12.4, H3ax-Neu5Ac) ppm.

$^{13}C$ NMR (151 MHz, $D_2O$): d = 102.9, 102.8, 102.6, 101.3, 100.5, 99.5, 97.0, 82.0, 80.4, 79.7, 78.7, 78.4, 78.1, 76.4, 76.2, 75.4, 75.3, 75.2, 74.4, 74.2, 73.5, 72.8, 72.3, 72.2, 72.0, 71.7, 71.0, 70.1, 69.4, 69.3, 68.6, 68.3, 67.6, 65.9, 62.3, 61.8, 61.5, 61.0, 60.0, 55.7, 55.0, 53.6, 51.7, 50.9, 39.6, 35.0, 22.4, 22.0 ppm.

ESI TOF-MS: for $C_{85}H_{139}N_8O_{61}$: m/z $[M-2H]^{2-}$ ; calcd: 1126.4077; found: 1126.4074.

**Preparation of compound 26**

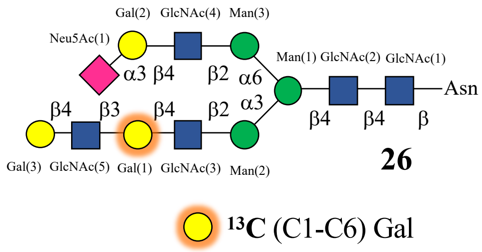


**26**

Compound **26** was prepared from **25** (2.6 mg, 1.2 µmol) using the general procedure for the installation of β1,3 Gal using B4GalT1. After P2 column purification, compound **26** was obtained as a white solid (2.6 mg, 92%).

$^1H$ NMR (600 MHz, $D_2O$): d = 5.03 (d, J < 2.0 Hz, 1H, H1-Man(2)), 4.99 (d, J = 9.8 Hz, 1H, H1-GlcNAc(1)), 4.83 (d, J < 2.0 Hz, 1H, H1-Man(3)), 4.69 (d, J = 2.6 Hz, 1H, H1-Man(1)), 4.59 (d, J = 9.7 Hz, 1H, H1-GlcNAc(5)), 4,53 (d, J = 7.6 Hz, 1H, H1-GlcNAc(2)), 4.47 (d, J = 7.8 Hz, 2H, H1-GlcNAc(3), H1-GlcNAc(4)), 4.43 (d, J = 7.7 Hz, 1H, H1-Gal(2)), 4.38 (d, J = 7.6 Hz, 2H, H1-Gal(1), H1-Gal(3)), 4.17 (dd, J = 2.6 Hz, J = 6.5 Hz, 1H, H2-Man(1)), 4.11 (dd, J < 2.0 Hz, J = 2.6 Hz, H2-Man(2)), 4.06 (dd, J < 2.0 Hz, J = 2.6 Hz, H4-Gal(1)), 4.00 (dd, J < 2.0 Hz, J = 2.6 Hz, H2-Man(3)), 3.99 (dd, J < 2.0 Hz, J = 2.5 Hz, H4-Gal(2)), 3.97-3.30 (m, 71H, H2-H6-GlcNAc(1), H2-H6-GlcNAc(2), H3-H6-Man(1), H3-H6-Man(2), H3-H6-Man(3), H2-H6-GlcNAc(4), H2-H6-GlcNAc(3), H2-H3, H5-H6-Gal(2), H2-H6 -Gal(1), H2-H6-GlcNAc(5), H2-H6 -Gal(3), H4-H9 Neu5Ac, αCH-Asn), 2.84-2.78 (m, 2H, βCH2-Asn), 2.64 (dd, J = 12.4, J = 4.6, H3eq-Neu5Ac), 1.99-1.95 (m, 18H, NHAc-GlcNAc(1), NHAc-GlcNAc(2), NHAc-GlcNAc(3), NHAc-GlcNAc(4), NHAc-GlcNAc(5), NHAc-Neu5Ac, 1.68 (t, J = 12.4, H3ax-Neu5Ac) ppm.

$^{13}C$ NMR (151 MHz, $D_2O$): d = 102.9, 102.8, 102.6, 101.3, 100.5, 99.5, 97.0, 82.0, 80.4, 79.7, 78.7, 78.4, 78.1, 77.8, 76.4, 76.2, 75.4, 75.3, 75.2, 74.4, 74.3, 74.2, 73.5, 72.8, 72.3, 72.2, 72.0, 71.7, 71.0, 70.1, 69.4, 69.3, 68.6, 68.3, 67.6, 65.9, 62.3, 61.8, 61.5, 61.0, 60.0, 59.7, 55.7, 55.0, 53.6, 51.7, 50.9, 39.6, 35.0, 22.4, 22.0 ppm.

ESI TOF-MS: for $C_{91}H_{163}N_8O_{66}$: m/z $[M-2H]^{2-}$ ; calcd: 1207.4341; found: 1207.4311.

**Preparation of compound 27**

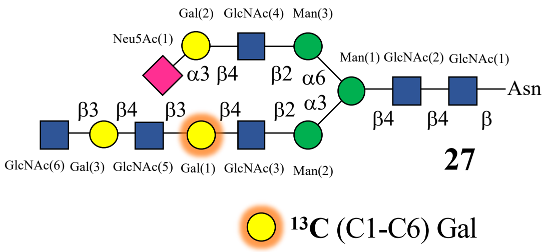


**27**

Compound **27** was prepared from **26** (2.6 mg, 1.1 µmol) using the general procedure for the installation of β1,4-GlcNAc using B3GnT2. After P2 column purification, compound **27** was obtained as a white solid (2.4 mg, 82%).

$^1H$ NMR (600 MHz, $D_2O$): d = 5.03 (d, J < 2.0 Hz, 1H, H1-Man(2)), 4.99 (d, J = 9.8 Hz, 1H, H1-GlcNAc(1)), 4.83 (d, J < 2.0 Hz, 1H, H1-Man(3)), 4.69 (d, J = 2.6 Hz, 1H, H1-Man(1)), 4.59 (d, J = 9.7 Hz, 2H, H1-GlcNAc(5), H1-GlcNAc(6)), 4,53 (d, J = 7.6 Hz,

1H, H1-GlcNAc(2)), 4.47 (d, J = 7.8 Hz, 2H, H1-GlcNAc(3), H1-GlcNAc(4)), 4.43 (d, J = 7.7 Hz, 1H, H1-Gal(2)), 4.38 (d, J = 7.6 Hz, 2H, H1-Gal(1), H1-Gal(3)), 4.17 (dd, J = 2.6 Hz, J = 6.5 Hz, 1H, H2-Man(1)), 4.11 (dd, J < 2.0 Hz, J = 2.6 Hz, H2-Man(2)), 4.06 (dd, J < 2.0 Hz, J = 2.6 Hz, 2H, H4-Gal(1), H4-Gal(3)), 4.00 (dd, J < 2.0 Hz, J = 2.6 Hz, H2-Man(3)), 3.99 (dd, J < 2.0 Hz, J = 2.5 Hz, H4-Gal(2)), 3.97-3.30 (m, 76H, H2-H6-GlcNAc(1), H2-H6-GlcNAc(2), H3-H6-Man(1), H3-H6-Man(2), H3-H6-Man(3), H2-H6-GlcNAc(4), H2-H6-GlcNAc(3), H2-H3, H5-H6-Gal(2), H2-H6 -Gal(1), H2-H6-GlcNAc(5), H2-H6 -Gal(3), H2-H6 GlcNAc(6), H4-H9 Neu5Ac, αCH-Asn), 2.84-2.78 (m, 2H, βCH2-Asn), 2.64 (dd, J = 12.4, J = 4.6, H3eq-Neu5Ac), 1.99-1.95 (m, 21H, NHAc-GlcNAc(1), NHAc-GlcNAc(2), NHAc-GlcNAc(3), NHAc-GlcNAc(4), NHAc-GlcNAc(5), NHAc-GlcNAc(6), NHAc-Neu5Ac, 1.68 (t, J = 12.4, H3ax-Neu5Ac) ppm.

$^{13}$C NMR (151 MHz, $D_2O$): d = 102.9, 102.8, 102.6, 101.3, 100.5, 99.5, 97.0, 82.0, 80.4, 79.7, 78.7, 78.4, 78.1, 77.8, 76.4, 76.2, 75.4, 75.3, 75.2, 74.4, 74.3, 74.2, 73.5, 72.8, 72.3, 72.2, 72.0, 71.7, 71.0, 70.1, 69.4, 69.3, 68.6, 68.3, 67.6, 65.9, 62.3, 61.8, 61.5, 61.0, 60.0, 59.7, 55.7, 55.0, 53.6, 51.7, 50.9, 39.6, 35.0, 22.4, 22.0 ppm.

ESI TOF-MS: for $C_{99}H_{162}N_9O_{71}$: m/z $[M-2H]^{2-}$ ; calcd: 1309.4755; found: 1309.4721.

**Preparation of compound 28**

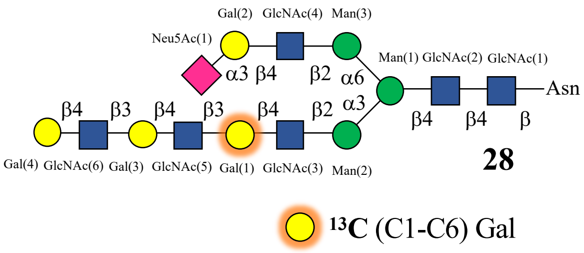


**28**

**28** was prepared from **27** (2.4 mg, 0.9 µmol) using the general procedure for the installation of β1,3-Gal using B4GalT1. After P2 column purification, compound **28** was obtained as a white solid (2.2 mg, 89%).

$^{1}$H NMR (600 MHz, $D_2O$): d = 5.03 (d, J < 2.0 Hz, 1H, H1-Man(2)), 4.99 (d, J = 9.8 Hz, 1H, H1-GlcNAc(1)), 4.83 (d, J < 2.0 Hz, 1H, H1-Man(3)), 4.69 (d, J = 2.6 Hz, 1H, H1-Man(1)), 4.59 (d, J = 9.7 Hz, 2H, H1-GlcNAc(5), H1-GlcNAc(6)), 4,53 (d, J = 7.6 Hz, 1H, H1-GlcNAc(2)), 4.47 (d, J = 7.8 Hz, 2H, H1-GlcNAc(3), H1-GlcNAc(4)), 4.43 (d, J = 7.7 Hz, 1H, H1-Gal(2)), 4.38 (d, J = 7.6 Hz, 3H, H1-Gal(1), H1-Gal(3), H1-Gal(4)), 4.17 (dd, J = 2.6 Hz, J = 6.5 Hz, 1H, H2-Man(1)), 4.11 (dd, J < 2.0 Hz, J = 2.6 Hz, H2-Man(2)), 4.06 (dd, J < 2.0 Hz, J = 2.6 Hz, 2H, H4-Gal(1), H4-Gal(3)), 4.00 (dd, J < 2.0 Hz, J = 2.6 Hz, H2-Man(3)), 3.99 (dd, J < 2.0 Hz, J = 2.5 Hz, H4-Gal(2)), 3.97-3.30 (m, 82H, H2-H6-GlcNAc(1), H2-H6-GlcNAc(2), H3-H6-Man(1), H3-H6-Man(2), H3-H6-Man(3), H2-H6-GlcNAc(4), H2-H6-GlcNAc(3), H2-H3, H5-H6-Gal(2), H2-H6-Gal(1), H2-H6-GlcNAc(5), H2-H6 -Gal(3), H2-H6 GlcNAc(6), H4-H9 Neu5Ac, H2-H6-Gal(4) αCH-Asn), 2.84-2.78 (m, 2H, βCH2-Asn), 2.64 (dd, J = 12.4, J = 4.6, H3eq-Neu5Ac), 1.99-1.95 (m, 21H, NHAc-GlcNAc(1), NHAc-GlcNAc(2), NHAc-GlcNAc(3), NHAc-GlcNAc(4), NHAc-GlcNAc(5), NHAc-GlcNAc(6), NHAc-Neu5Ac, 1.68 (t, J = 12.4, H3ax-Neu5Ac) ppm.

$^{13}$C NMR (151 MHz, $D_2O$): d = 102.9, 102.8, 102.6, 101.3, 100.5, 99.5, 97.0, 82.0, 80.4, 79.7, 78.7, 78.4, 78.1, 77.8, 76.4, 76.2, 75.4, 75.3, 75.2, 74.4, 74.3, 74.2, 73.5, 72.8, 72.3, 72.2, 72.0, 71.7, 71.0, 70.1, 69.4, 69.3, 68.6, 68.3, 67.6, 65.9, 62.3, 61.8, 61.5, 61.0, 60.0, 59.7, 55.7, 55.0, 53.6, 51.7, 50.9, 39.6, 35.0, 22.4, 22.0 ppm.

ESI TOF-MS: for $C_{105}H_{173}N_9O_{76}$: m/z $[M-3H]^{3-}$ ; calcd: 926.6655; found: 926.6622.

**Preparation of compound 29**

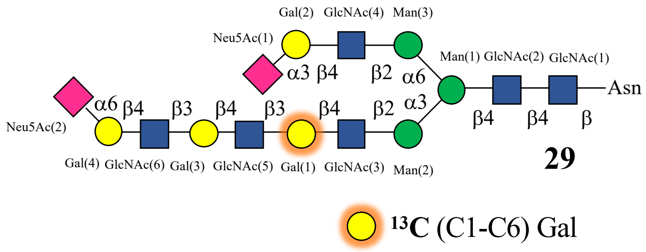


**29**

Compound **29** was prepared from **28** (2.2 mg, 0.8 µmol) using the general procedure for the installation of α2,6-Neu5Ac using ST6Gal1. After P2 column purification, compound **29** was obtained as a white solid (2.1 mg, 86%).

$^1$H NMR (600 MHz, $D_2O$): d = 5.03 (d, J < 2.0 Hz, 1H, H1-Man(2)), 4.99 (d, J = 9.8 Hz, 1H, H1-GlcNAc(1)), 4.83 (d, J < 2.0 Hz, 1H, H1-Man(3)), 4.69 (d, J = 2.6 Hz, 1H, H1-Man(1)), 4.59 (d, J = 9.7 Hz, 2H, H1-GlcNAc(5), H1-GlcNAc(6)), 4,53 (d, J = 7.6 Hz, 1H, H1-GlcNAc(2)), 4.47 (d, J = 7.8 Hz, 2H, H1-GlcNAc(3), H1-GlcNAc(4)), 4.43 (d, J = 7.7 Hz, 1H, H1-Gal(2)), 4.38 (d, J = 7.6 Hz, 2H, H1-Gal(1), H1-Gal(3)), 4.35 (d, J = 7.8 Hz, 1H, H1-Gal(4)), 4.17 (dd, J = 2.6 Hz, J = 6.5 Hz, 1H, H2-Man(1)), 4.11 (dd, J < 2.0 Hz, J = 2.6 Hz, H2-Man(2)), 4.06 (dd, J < 2.0 Hz, J = 2.6 Hz, 3H, H4-Gal(1), H4-Gal(3), H4-Gal(4)), 4.00 (dd, J < 2.0 Hz, J = 2.6 Hz, H2-Man(3)), 3.99 (dd, J < 2.0 Hz, J = 2.5 Hz, H4-Gal(2)), 3.97-3.30 (m, 94H, H2-H6-GlcNAc(1), H2-H6-GlcNAc(2), H3-H6-Man(1), H3-H6-Man(2), H3-H6-Man(3), H2-H6-GlcNAc(4), H2-H6-GlcNAc(3), H2-H3, H5-H6-Gal(2), H2-H6-Gal(1), H2-H6-GlcNAc(5), H2-H6 -Gal(3), H2-H6 GlcNAc(6), H4-H9 Neu5Ac(1), H4-H9 Neu5Ac(2), H2-H3, H5-H6-Gal(4) αCH-Asn), 2.84-2.78 (m, 2H, βCH2-Asn), 2.64 (dd, J = 12.4, J = 4.6, H3eq-Neu5Ac(1), 2.54 (dd, J = 12.4, J = 4.6, H3eq-Neu5Ac(2), 1.99-1.95 (m, 24H, NHAc-GlcNAc(1), NHAc-GlcNAc(2), NHAc-GlcNAc(3), NHAc-GlcNAc(4), NHAc-GlcNAc(5), NHAc-GlcNAc(6), NHAc-Neu5Ac(1), NHAc-Neu5Ac(2), 1.68 (t, J = 12.4, H3ax-Neu5Ac(1), 1.58 (t, J = 12.4, H3ax-Neu5Ac(2) ppm.

$^{13}$C NMR (151 MHz, $D_2O$): d = 102.9, 102.8, 102.6, 101.3, 100.5, 99.5, 97.0, 82.0, 80.4, 79.7, 78.7, 78.4, 78.1, 77.8, 76.4, 76.2, 75.4, 75.3, 75.2, 74.4, 74.3, 74.2, 73.5, 72.8, 72.3, 72.2, 72.0, 71.7, 71.0, 70.1, 69.4, 69.3, 68.6, 68.3, 67.6, 67.2, 65.9, 62.3, 61.8, 61.6, 61.5, 61.0, 60.0, 59.7, 55.7, 55.0, 53.6, 51.7, 50.9, 50.8, 39.6, 39.1, 35.0, 22.4, 22.0 ppm.

ESI TOF-MS: for $C_{116}H_{190}N_{10}O_{84}$: m/z $[M-3H]^{3-}$ ; calcd: 1023.6973; found: 1023.6954.

**Preparation and characterization of compound 2, 3, 4 and 5**

Compound **3** was obtained from **16** (0.7 mg, 0.25 µmol) by using α1,2-fucosidase for the hydrolysis of the α1,2 Fuc. After P2 column purification, **3** was obtained as a white solid (0.6 mg, 96%).

Compound **4** was obtained from **21** (0.55 mg, 0.20 µmol) by using α1,2-fucosidase for the hydrolysis of the α1,2 Fuc. After P2 column purification, **4** was obtained as a white solid (0.45 mg, 90%).

Compound **5** was obtained from **29** (2.0 mg, 0.70 µmol) by selective hydrolysis of the α2,3-Neu5Ac. After P2 column purification, **5** was obtained as a white solid (1.45 mg, 74%).

NMR Nomenclature

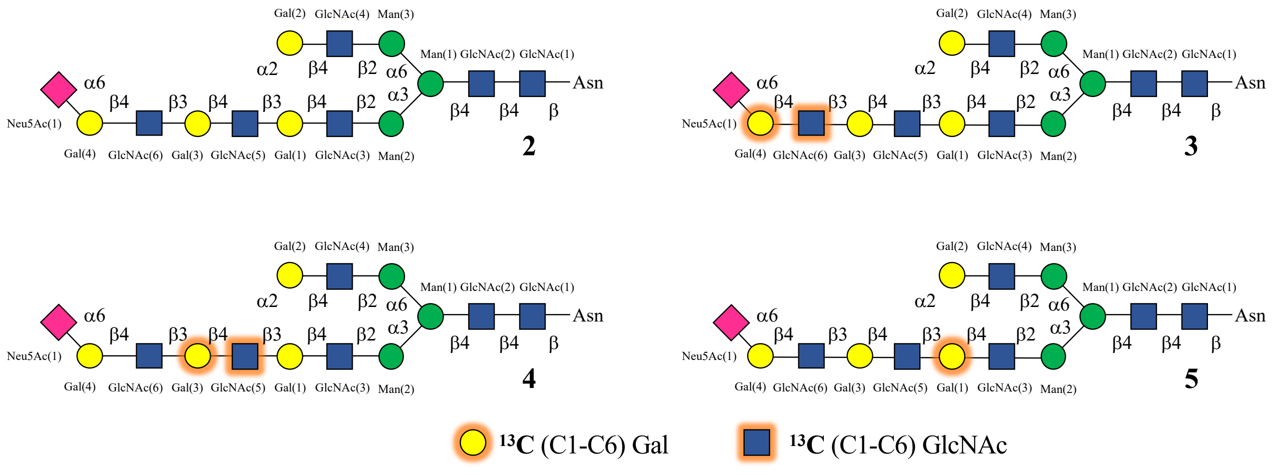


$^{13}C$; $^{1}H$ (600 MHz; $D_2O$). δ (ppm).

| | C1-H1 | C2-H2 | C3;H3 | C4;H4 | C5-H5 | C6-H6R;H6S | C7-H7 | C8-H8 | C9-H9 | NHAc | Cα-Hα | Cβ-Hβ |
|---|---|---|---|---|---|---|---|---|---|---|---|---|
| **GlcNAc(1)** | 78.1; 4.99 | 53.6; 3.78 | 72.2; 3.67 | 78.7; 3.58 | 76.2; 3.50 | 60.0; 3.56; 3.75 | -[a] | -[a] | -[a] | 22.4-22.0; 1.99-1.93 21H | -[a] | -[a] |
| **GlcNAc(2)** | 101.3; 4.53 | 55.0; 3.71 | 73.5; 3.66 | 79.7; 3.65 | 74.4; 3.51 | 60.0; 3.56; 3.75 | -[a] | -[a] | -[a] | 22.4-22.0; 1.99-1.93 21H | -[a] | -[a] |
| **Man(1)** | 100.5; 4.69 | 70.1; 4.17 | 80.4; 3.69 | 65.9; 3.69 | 74.2; 3.56 | 65.9; 3.86; 3.71 | -[a] | -[a] | -[a] | -[a] | -[a] | -[a] |
| **Man(2)** | 99.5; 5.03 | 76.4; 4.11 | 69.2; 3.81 | 67.3; 3.42 | 73.5; 3.66 | 61.5; 3.83; 3.53 | -[a] | -[a] | -[a] | -[a] | -[a] | -[a] |
| **GlcNAc(3)** | 99.5; 4.50 | 55.0; 3.67 | 75.3; 3.63 | 78.4; 3.65 | 74.8; 3.49 | 59.9; 3.89; 3.76 | -[a] | -[a] | -[a] | 22.4-22.0; 1.99-1.93 21H | -[a] | -[a] |
| **Gal(1)** | 102.8; 4.32 | 69.8; 3.44 | 81.9; 3.59 | 68.2; 4.02 | 74.7; 3.58 | 60.9; 3.63; 3.59 | -[a] | -[a] | -[a] | -[a] | -[a] | -[a] |
| **Man(3)** | 97.0; 4.83 | 76.4; 4.00 | 69.2; 3.81 | 67.3; 3.41 | 72.9; 3.53 | 61.8; 3.83; 3.54 | -[a] | -[a] | -[a] | -[a] | -[a] | -[a] |
| **GlcNAc(4)** | 99.5; 4.50 | 55.0; 3.67 | 75.3; 3.63 | 78.4; 3.65 | 74.8; 3.49 | 59.9; 3.89; 3.76 | -[a] | -[a] | -[a] | 22.4-22.0; 1.99-1.93 21H | -[a] | -[a] |
| **Gal(2)** | 102.9; 4.38 | 71.0; 3.45 | 72.3; 3.58 | 68.6; 3.84 | 72.0; 3.66 | 61.0; 3.66; 3.66 | -[a] | -[a] | -[a] | -[a] | -[a] | -[a] |
| **GlcNAc(5)** | 102.7; 4.56 | 55.0; 3.66 | 72.0; 3.59 | 77.9; 3.59 | 74.4; 3.45 | 59.72; 3.82; 3.71 | -[a] | -[a] | -[a] | 22.4-22.0; 1.99-1.93 21H | -[a] | -[a] |
| **Gal(3)** | 102.8; 4.33 | 69.9; 3.44 | 81.9; 3.60 | 68.2; 4.02 | 74.7; 3.58 | 60.9; 3.63; 3.59 | -[a] | -[a] | -[a] | -[a] | -[a] | -[a] |
| **GlcNAc(6)** | 102.8; 4.55 | 54.8; 3.66 | 71.1; 3.65 | 79.3; 3.53 | 72.6; 3.69 | 59.0; 3.81; 3.71 | -[a] | -[a] | -[a] | 22.4-22.0; 1.99-1.93 21H | -[a] | -[a] |
| **Gal(4)** | 102.4; 4.32 | 69.6; 3.30 | 71.3; 3.54 | 67.2; 3.79 | 72.6; 3.69 | 62.2; 3.86; 3.40 | -[a] | -[a] | -[a] | -[a] | -[a] | -[a] |

| | | | | | | | | | | | | |
|---|---|---|---|---|---|---|---|---|---|---|---|---|
| **Neu5Ac(1)** | -[a] | -[a] | 39.1; 2.54; 1.58 | 67.2; 3.52 | 50.8; 3.68 | 71.4; 3.57 | 67.3; 3.43 | 70.5; 3.76 | 61.6; 3.74; 3.51 | 22.4-22.0; 1.99-1.93 21H | -[a] | -[a] |
| **Asn** | -[a] | -[a] | -[a] | -[a] | -[a] | -[a] | -[a] | -[a] | -[a] | -[a] | 50.9; 3.90 | 35.0; 2.84; 2.78 |

[a]Not applicable

ESI TOF-MS: for $C_{105}H_{173}N_9O_{76}$: m/z $[M-3H]^{3-}$ ; (**2**) calcd: 924.6588; found: 924.6602.

(**3**) calcd: 928.6722; found: 928.6695. (**4**) calcd: 928.6722; found: 928.6702. (**5**) calcd: 926.6655; found: 926.6643.

### 1.1 NMR Spectra

Proton spectrum of **11**, 600 MHz, $D_2O$.

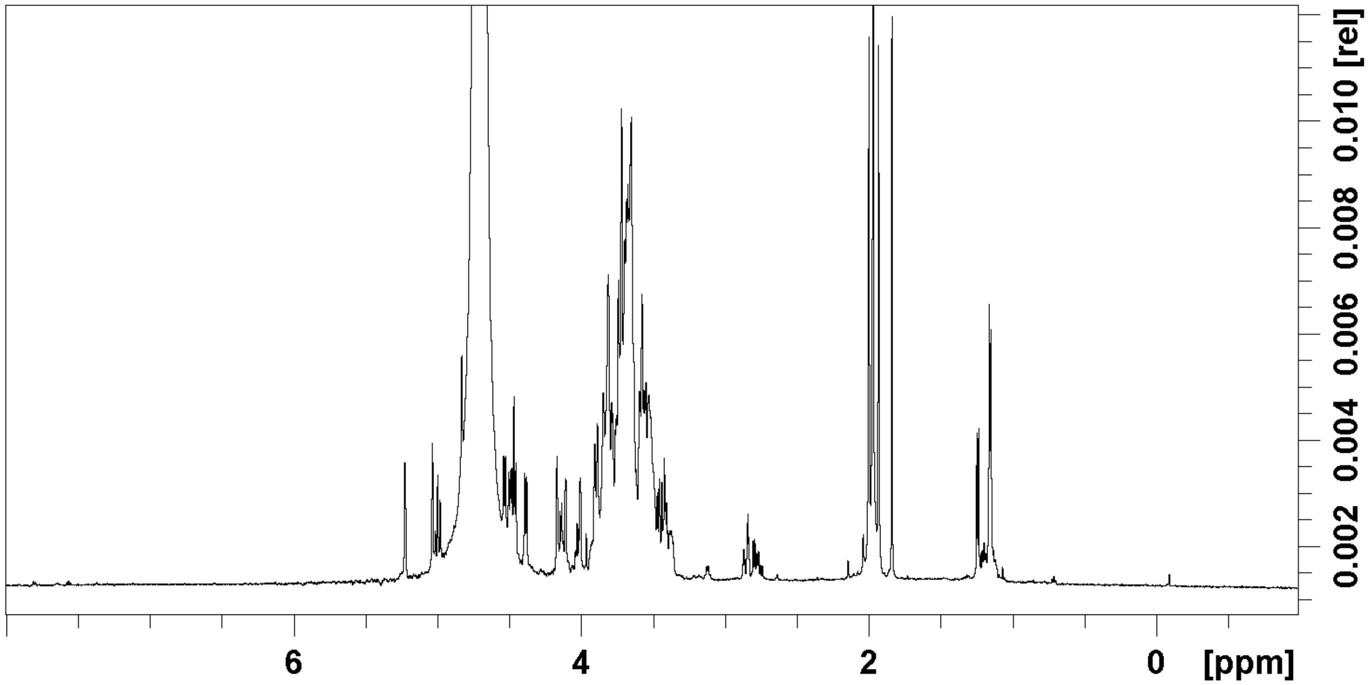


COSY spectrum of **11**, 600 MHz, $D_2O$.

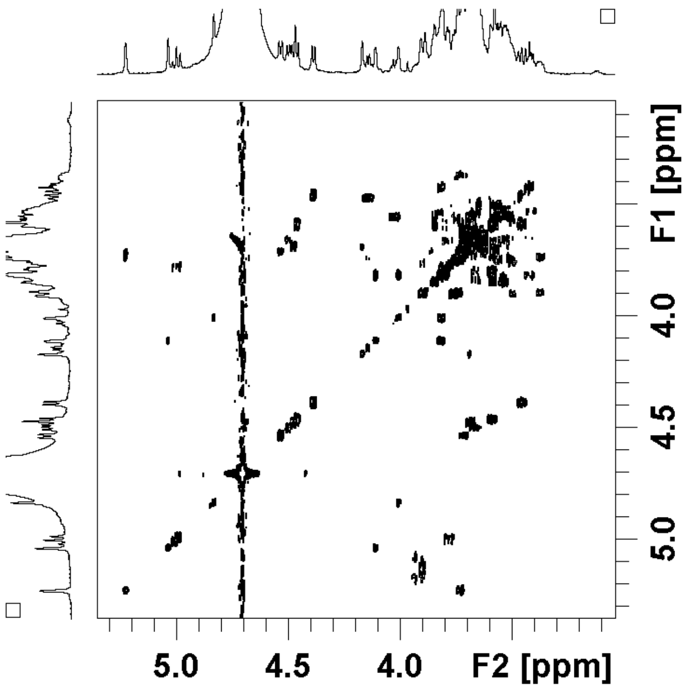


TOCSY spectrum of **11**, 600 MHz, $D_2O$.

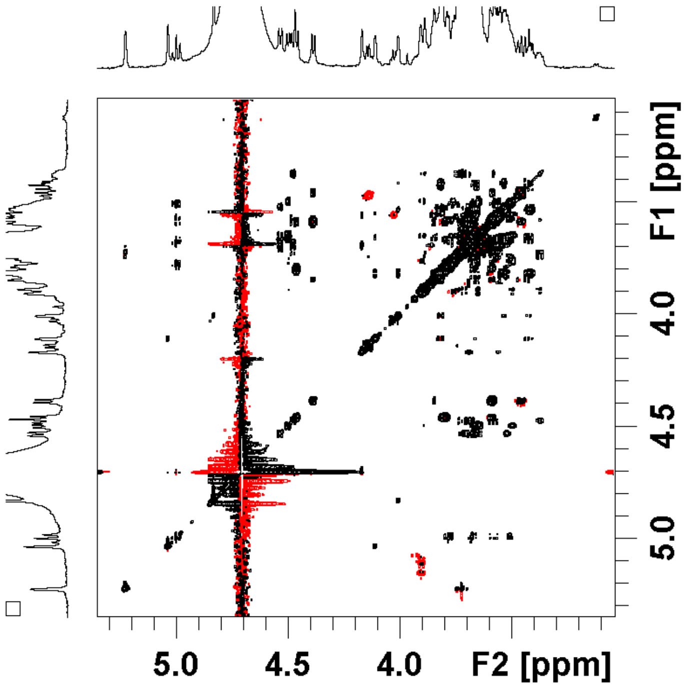


NOESY spectrum of **11**, 600 MHz, $D_2O$.

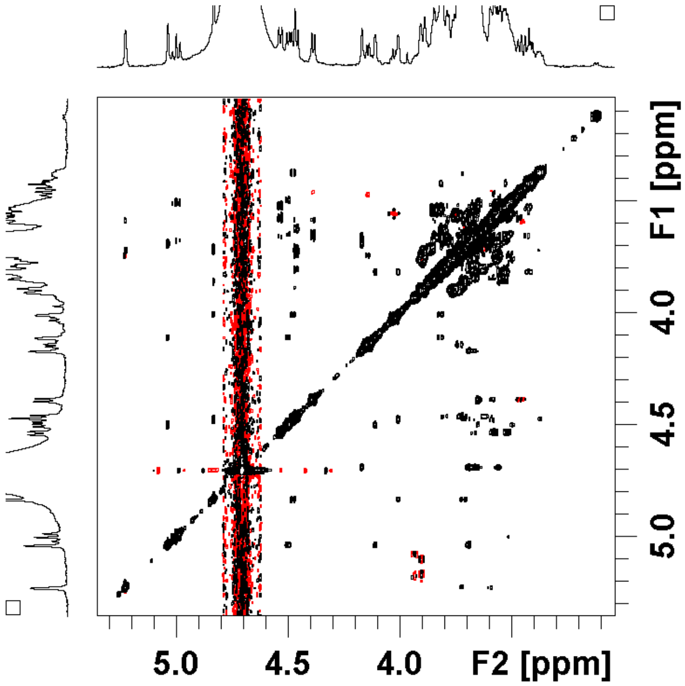


[1]H-[13]C-HSQC spectrum of **11**, 600 MHz, $D_2O$.

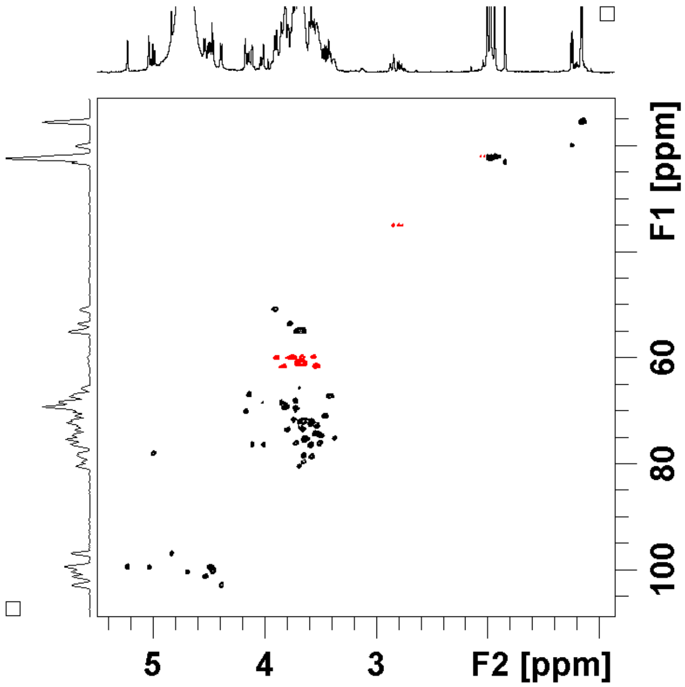


Proton spectrum of **17**, 600 MHz, $D_2O$.

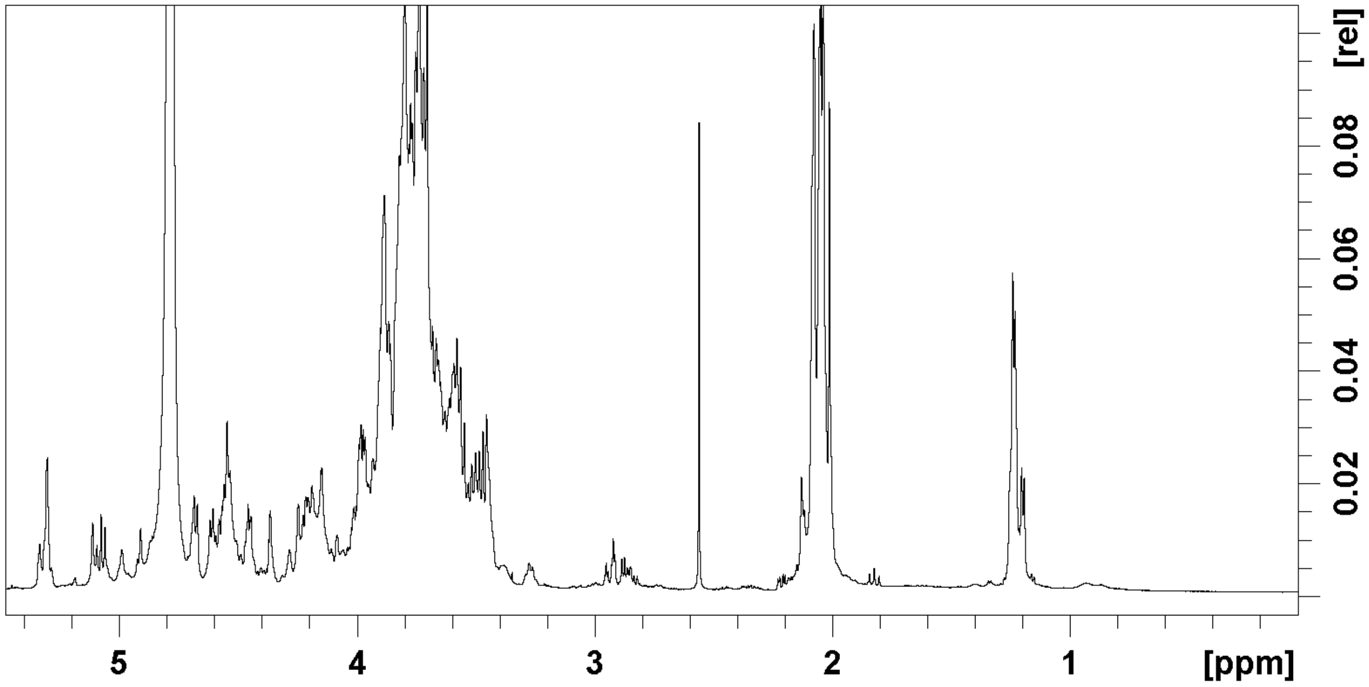


$^{1}H$-$^{13}C$-HSQC spectrum of **17**, 600 MHz, $D_2O$.

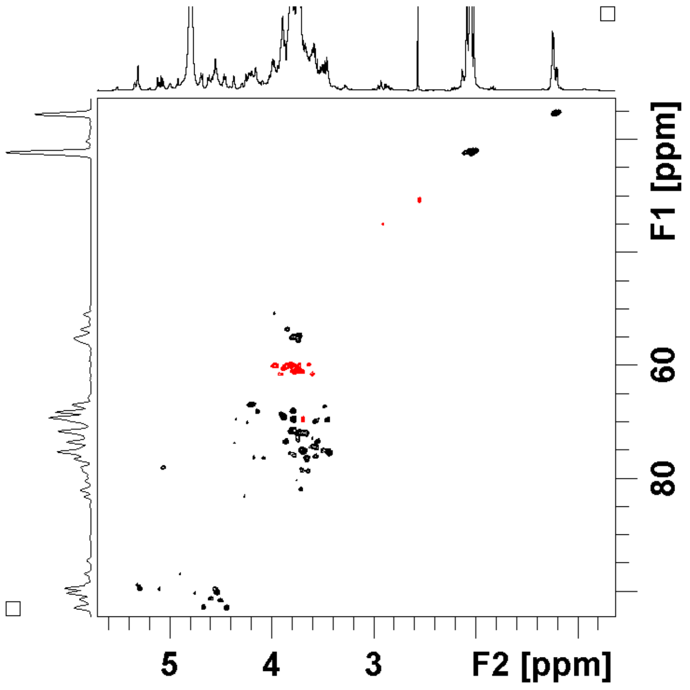


**Supplementary Fig. 1.** NMR spectra of compound **3**. a) full $^{1}$H-NMR spectrum in presence of TSP as internal reference. b) Zoom of the $^{1}$H-NMR spectrum. c) $^{1}$H-$^{13}$C HSQC spectrum showing all NMR signals (NS=128). d) $^{1}$H-$^{13}$C HSQC spectrum showing the NMR signals of the labelled residues, exclusively (NS=2).

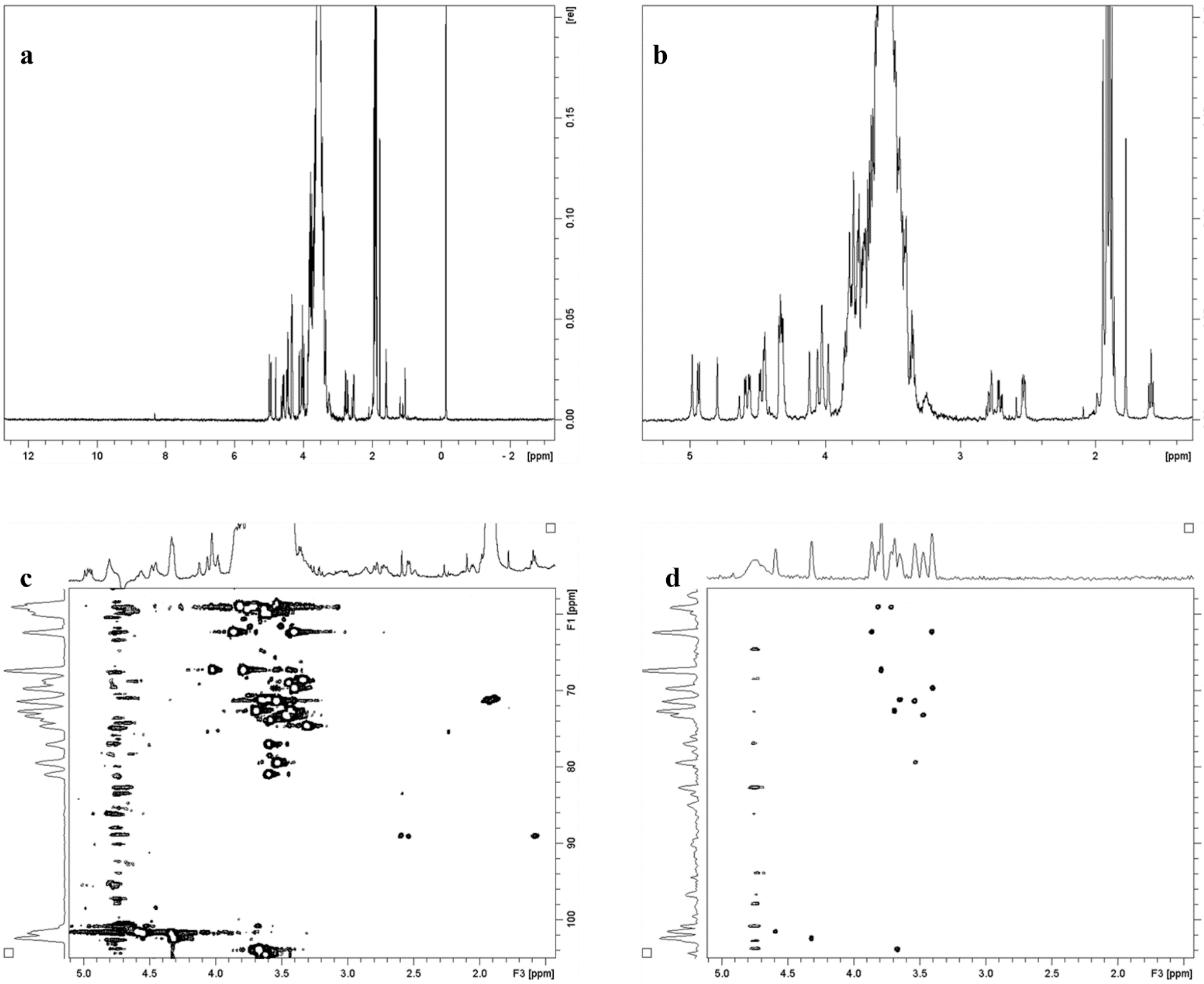
a
b
c
d
[rel]
[ppm]
F1 [ppm]
F3 [ppm]

**Supplementary Fig. 2.** NMR spectra of compound **4**. a) full $^{1}H$-NMR spectrum in presence of TSP as internal reference. b) Zoom of the $^{1}H$-NMR spectrum. c) $^{1}H$-$^{13}C$ HSQC spectrum showing all NMR signals (NS=128). d) $^{1}H$-$^{13}C$ HSQC spectrum showing the NMR signals of the labelled residues, exclusively (NS=2).

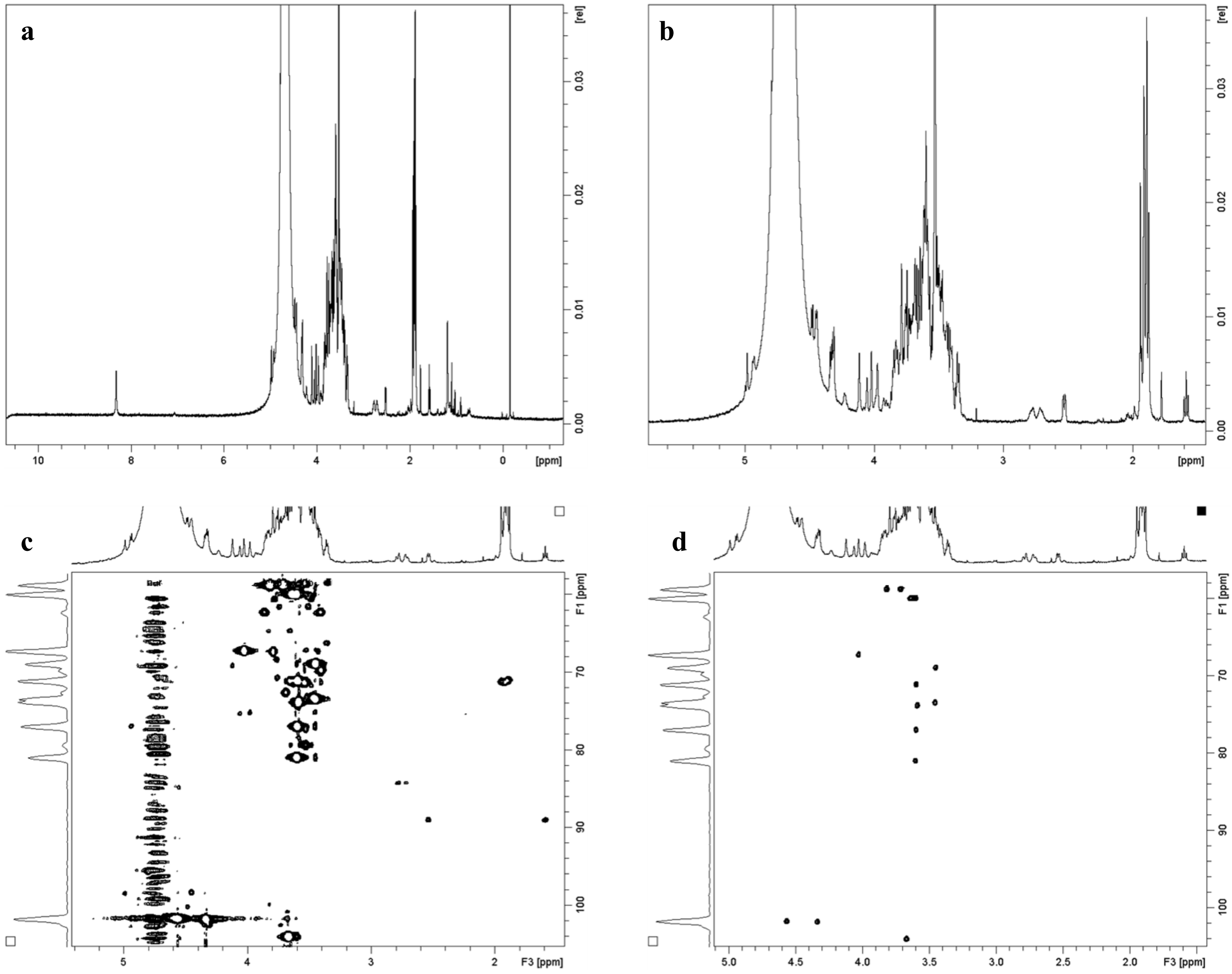


Proton spectrum of **28**, 600 MHz, $D_2O$.

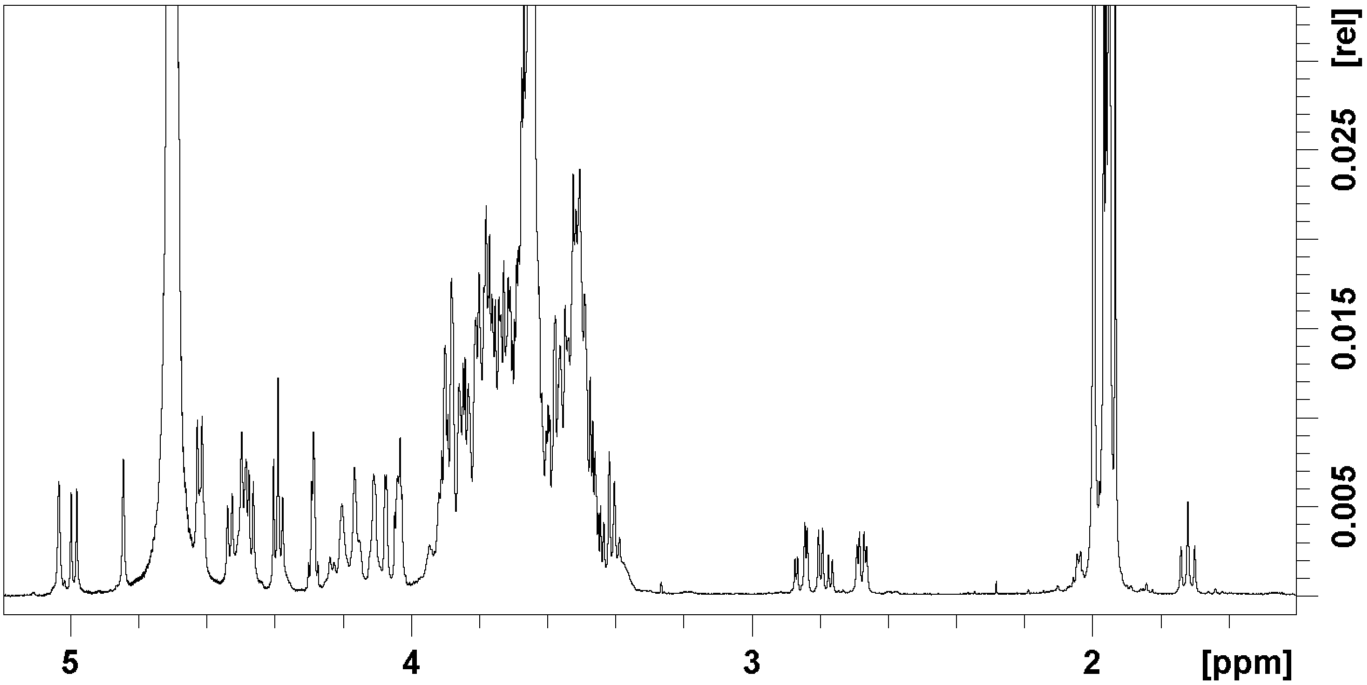


$^{1}H$-$^{13}C$-HSQC spectrum of **28**, 600 MHz, $D_2O$.

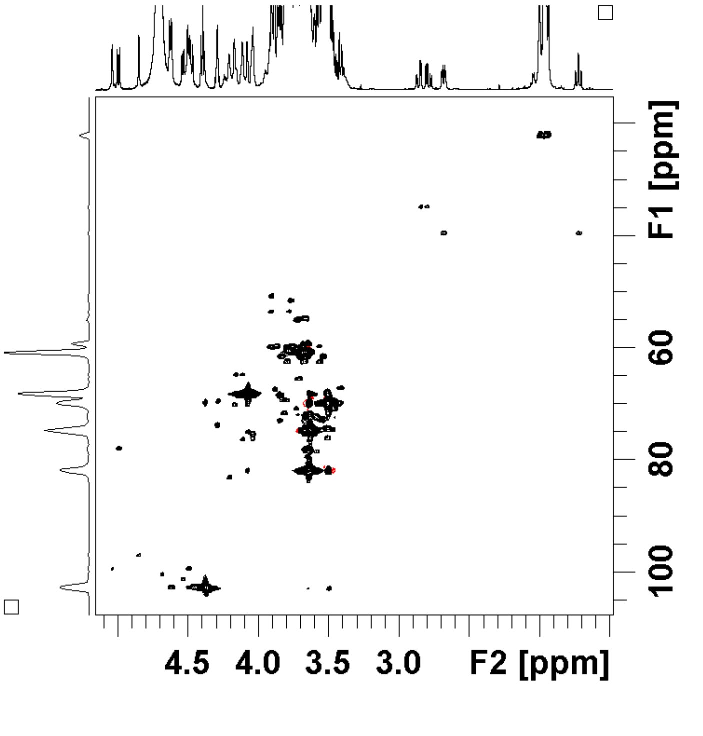


Proton spectrum of **29**, 600 MHz, $D_2O$.

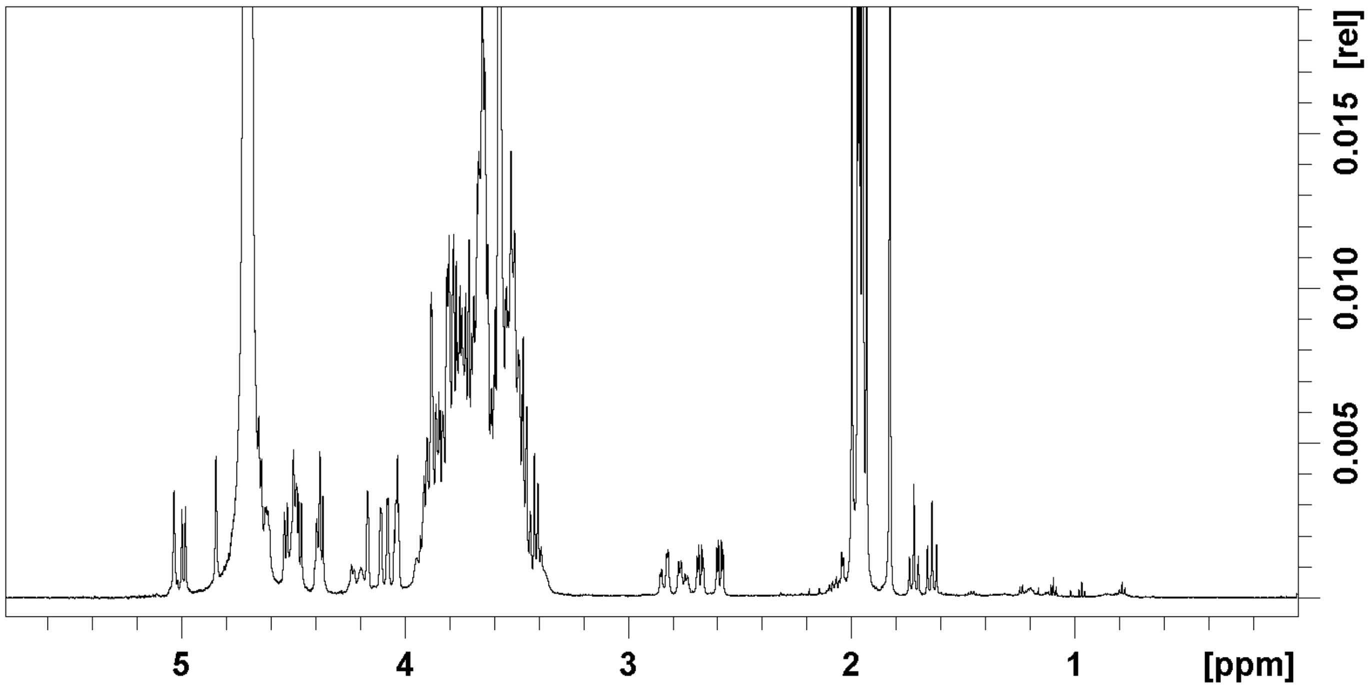


$^{1}H$-$^{13}C$-HSQC spectrum of **29**, 600 MHz, $D_2O$.

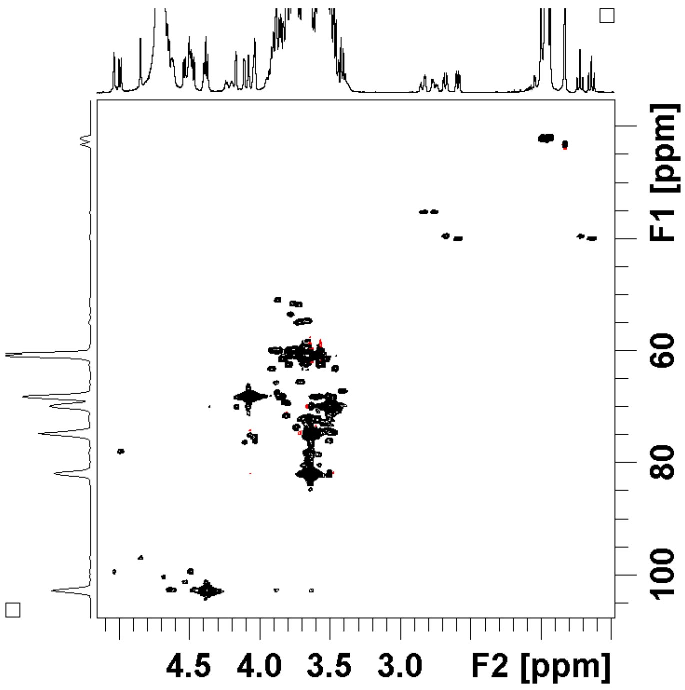

**Supplementary Fig. 3.** NMR spectra of compound **5**. a) full $^{1}$H-NMR spectrum in presence of TSP as internal reference. b) Zoom of the $^{1}$H-NMR spectrum. In a and b, 1H-NMR spectra (top) and 1H-{13C} decoupled spectra (bottom). c) $^{1}$H-$^{13}$C HSQC spectrum showing all NMR signals (NS=128). d) $^{1}$H-$^{13}$C HSQC spectrum showing the NMR signals of the labelled residues, exclusively (NS=2).

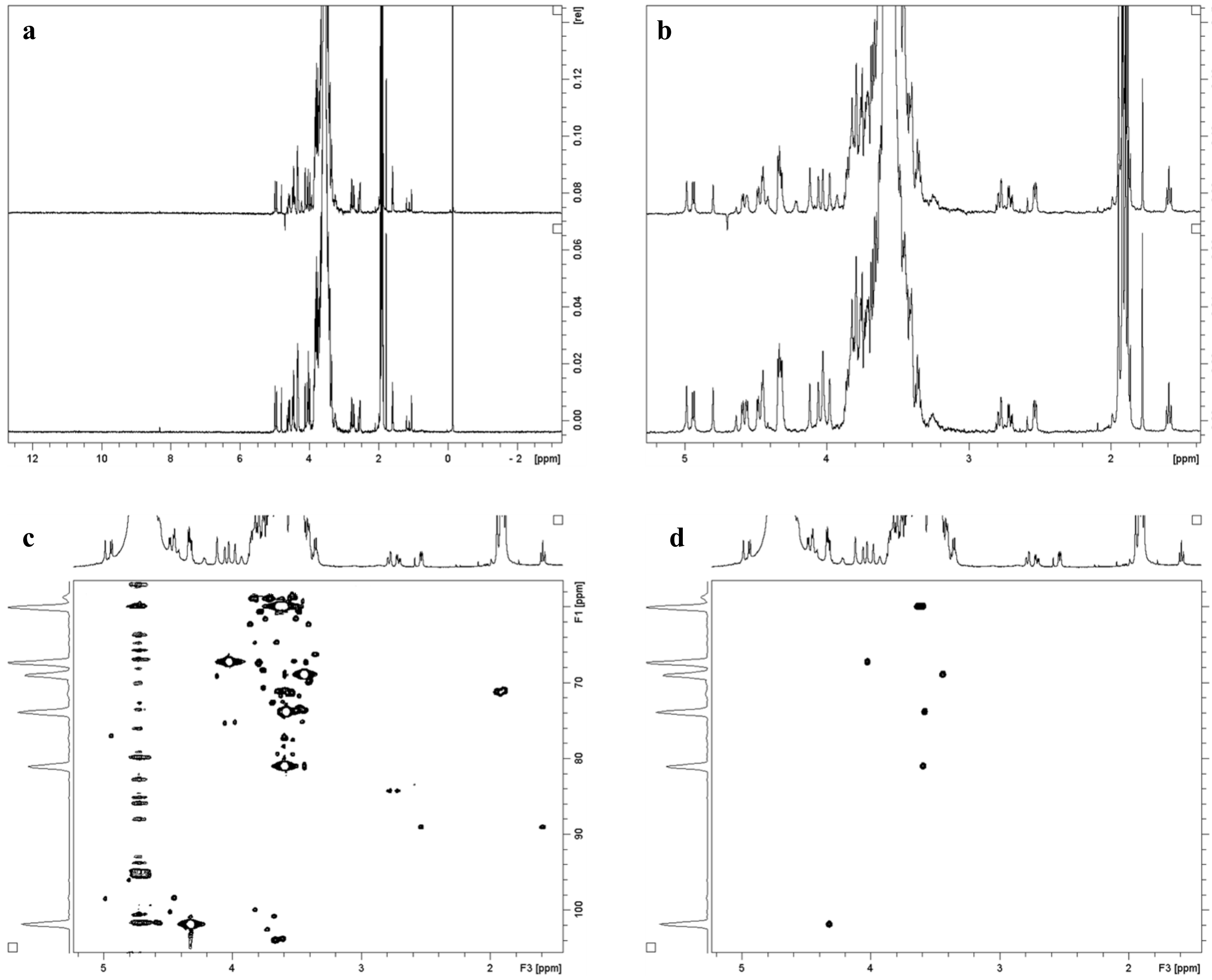

### 2. Synthesis of compound 6

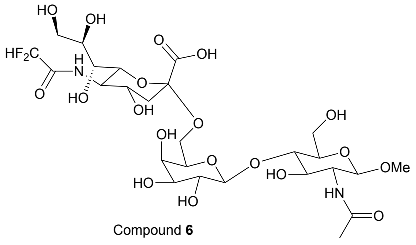


Compound **6**

Mannosamine hydrochloride (300 mg, 1.4 mmol) was dissolved in water/MeOH (1:1, 10 mL) and filtered with $HCO_3^-$ resin to release free mannosamine. 1 mL of triethylamine and 1 mlLof ethyl difluoroacetate (7 mmol) were added to the solution, which was then stirred at room temperature for 20 h. The reaction mixture was concentrated *in vacuo*. The residue was purified by silica gel chromatography to give the Man($CHF_2CONH$-). The Man($CHF_2CONH$-) (1.0 eq, 0.12 mmol, 18 mM) and sodium pyruvate (5.0 eq, 0.6 mmol, 90 mM) were dissolved in MiliQ water in a 50 mL centrifuge tube containing Tris-HCl buffer (100 mM, pH 7.5) and $MgCl_2$ (20 mM). After the addition of sialic acid aldolase, the reaction was incubated at 37 °C for 2 days with shaking. The progress of the reaction was monitored by TLC and LC-MS. Then to this mixture was added CTP (2.0 eq) and the pH was adjusted to 8.5 with 1M NaOH, after which NmCSS and PPA were added. The reaction was incubated at 37 °C for 6 h with shaking. Product formation was monitored by LC-MS. After the generation of CMP-Neu5($CHF_2CONH$) was completed, ST6Gal1 and LacNAc acceptor were added to the reaction mixture, which was incubated for 1 day at 37 °C with shaking and product formation was monitored by LC-MS. The reaction mixture was lyophilized and purified by size-exclusion column chromatography using P-2 BioGel eluting with a 50 mM ammonium bicarbonate solution to give **6** as white solid (289 mg, 30%).

$^1$H (600 MHz; $D_2O$). δ (ppm). 6.17 (t, $^2J_{H\text{-}F}$ = 54 Hz, $CHF_2$-), 4.50 (d, $^3J_{H\text{-}H}$ = 8 Hz, 1H), 4.45 (d, $^3J_{H\text{-}H}$ =7.8 Hz, 1H), 3.50-4.05 (m, 19H), 3.52 (s, 3H, OMe), 2.69 (dd, $^3J_{H\text{-}H}$ = 4.75 Hz, $^2J_{H\text{-}H}$ = 12.62 Hz, 1H, H-3eq), 2.08 (s, 3H, NHAc), 1.74 (t, $^3J_{H\text{-}H}$ = $^2J_{H\text{-}H}$ = 12.19 Hz, H-3ax).

ESI TOF-MS: for $C_{26}H_{42}F_2N_2O_{19}$: m/z [M-H]$^-$ ; calcd: 723.2277 ; found: 723.2546.

**3.1. Supplementary Fig. 4.** 2D STD-$^{1}$H,$^{13}$C-HSQC NMR experiments for the interaction of compound **4** with HA proteins. Top, the off-resonance spectrum and the corresponding STD-HSQC spectra for the three HA proteins in complex with compound **4**. Bottom, sum of STD effects for all signals of the Gal (orange) and the GlcNAc (cyan) of compounds **4** in complex with the three HA proteins.

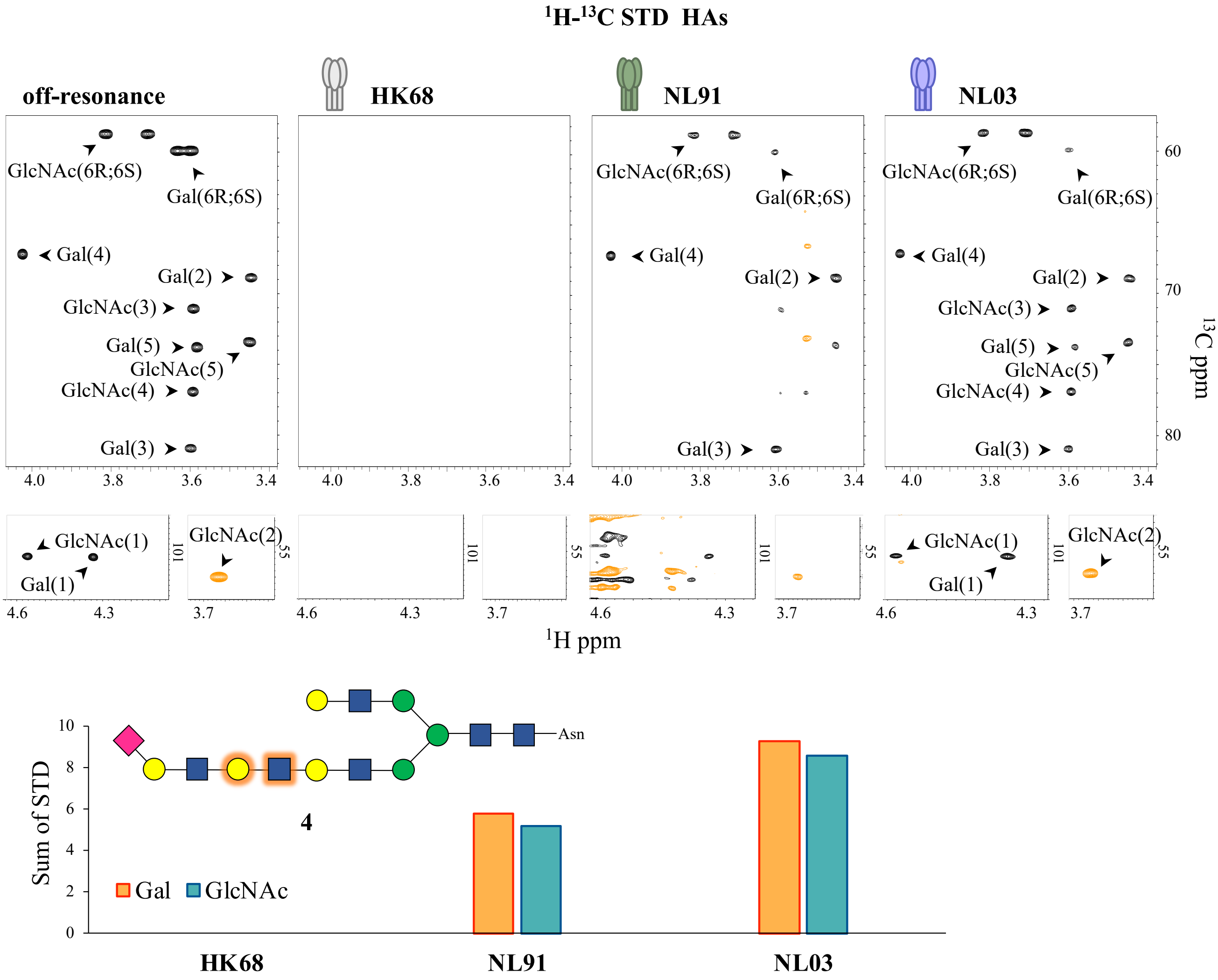

**3.2. Supplementary Fig. 5.** 2D STD-$^{1}$H,$^{13}$C-HSQC NMR experiments for the interaction of compound **5** with HA proteins. Top, the off-resonance spectrum and the corresponding STD-HSQC spectra for the three HA proteins in complex with compound **5**. Bottom, sum of STD effects for all signals of the Gal (orange) of compounds **5** in complex with the three HA proteins.

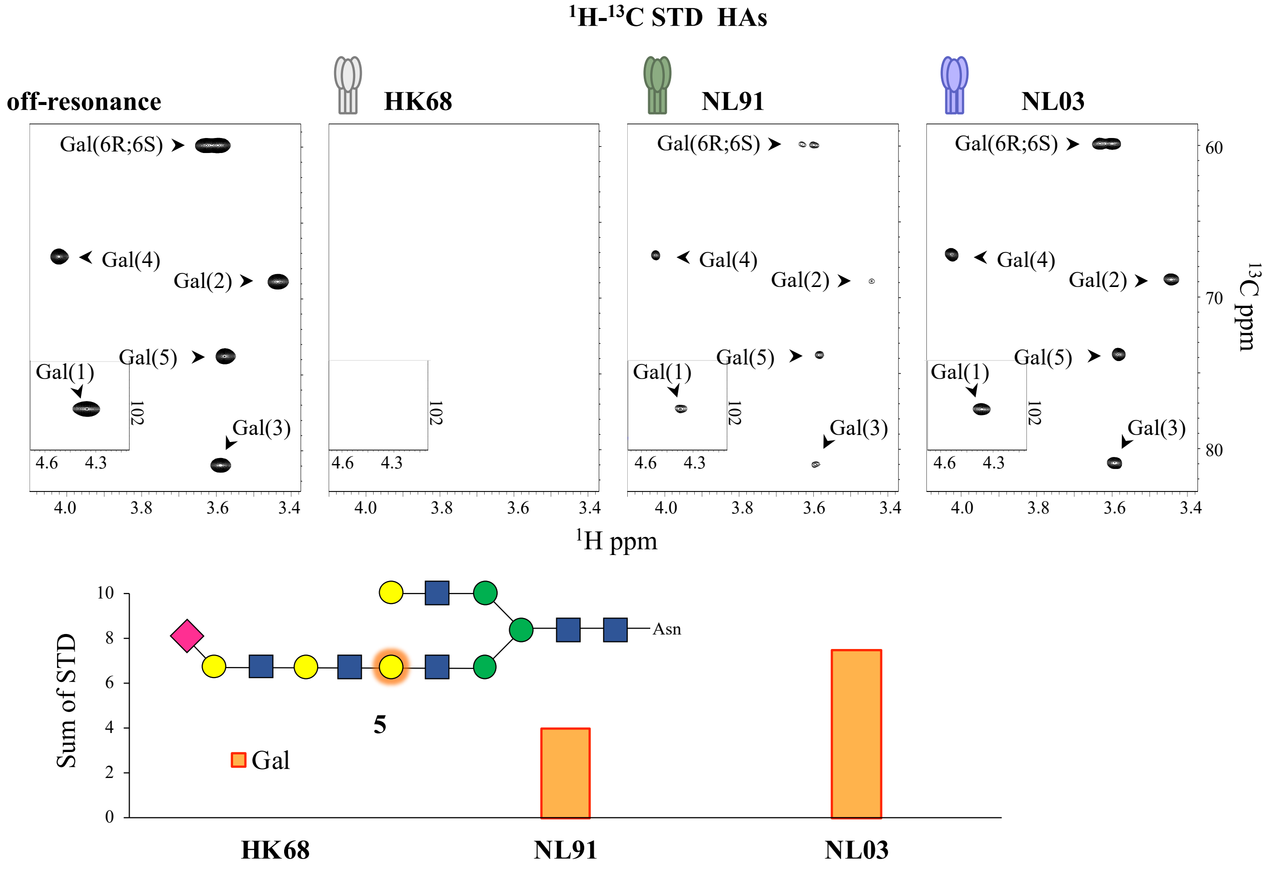

**4. Supplementary Fig. 6.** Details of 2D STD NMR results for the interaction of compound **5** with HA NL03 protein.

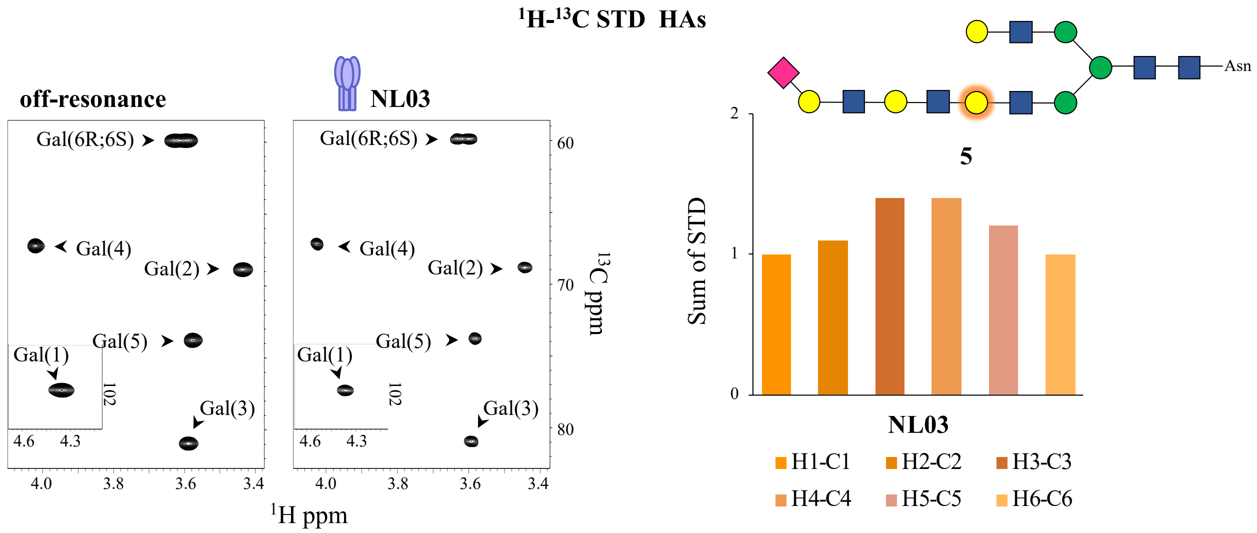

**5. Supplementary Fig. 7.** Details of the intermolecular interactions between **2** and the HAs proteins as derived by molecular modelling.

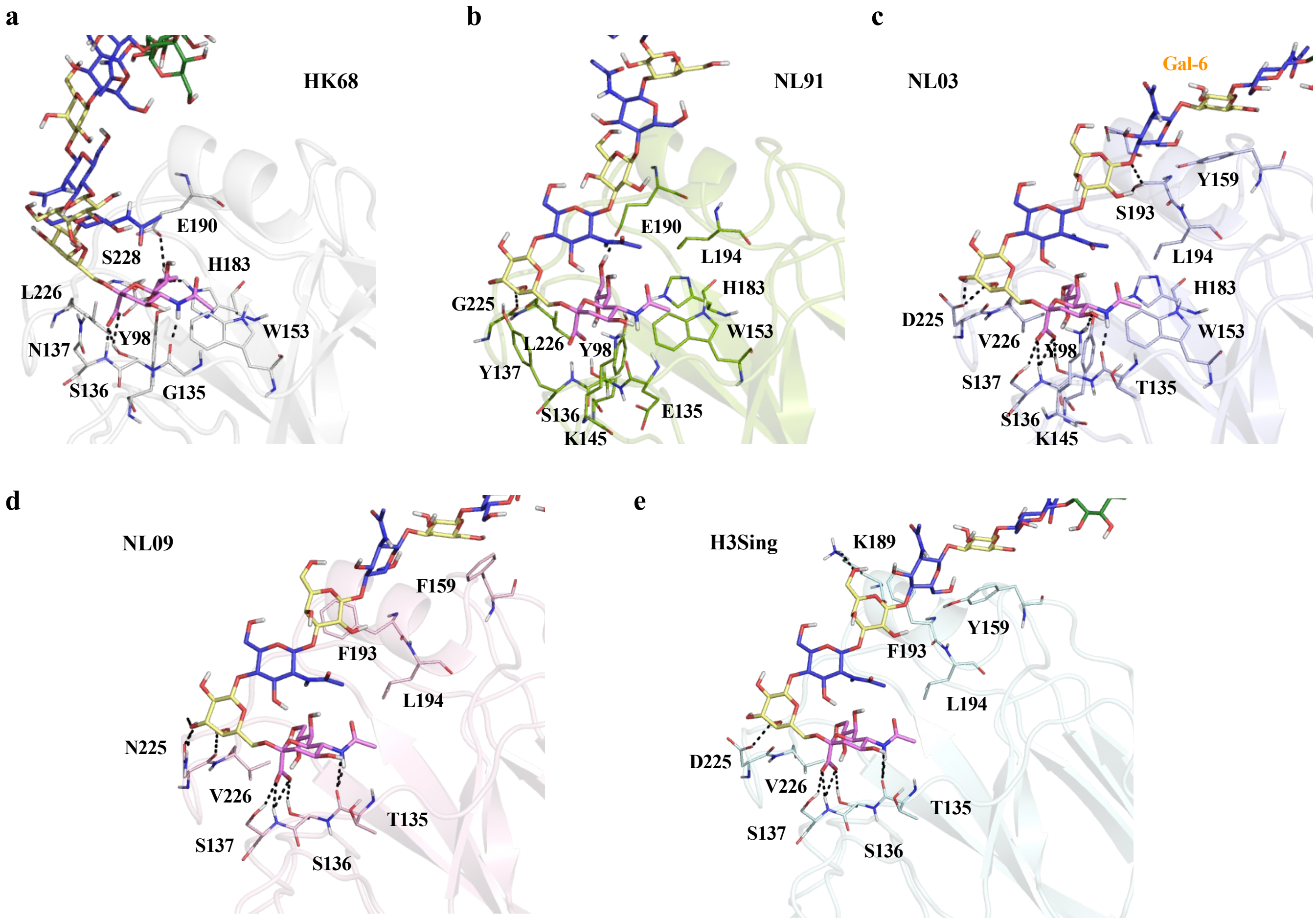

**6. Supplementary Fig. 8.** Snapshoot of the **2**/HK68 MD trajectory.

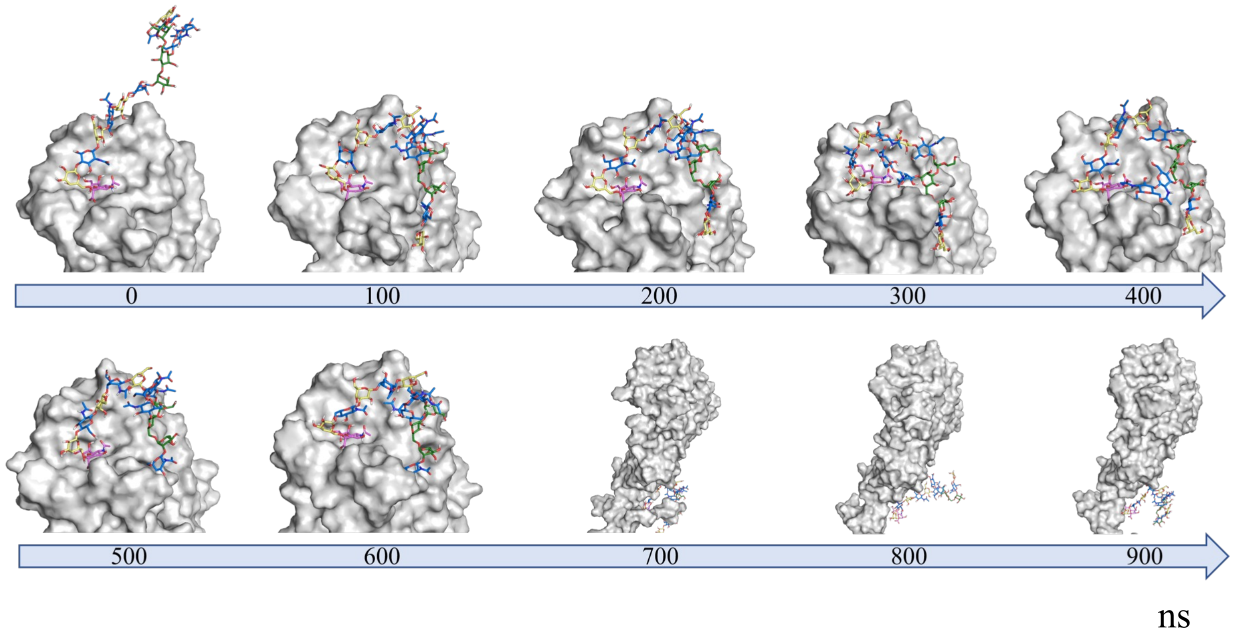


**7. Supplementary Fig. 9.** Snapshoot of the **2**/NL91 MD trajectory.

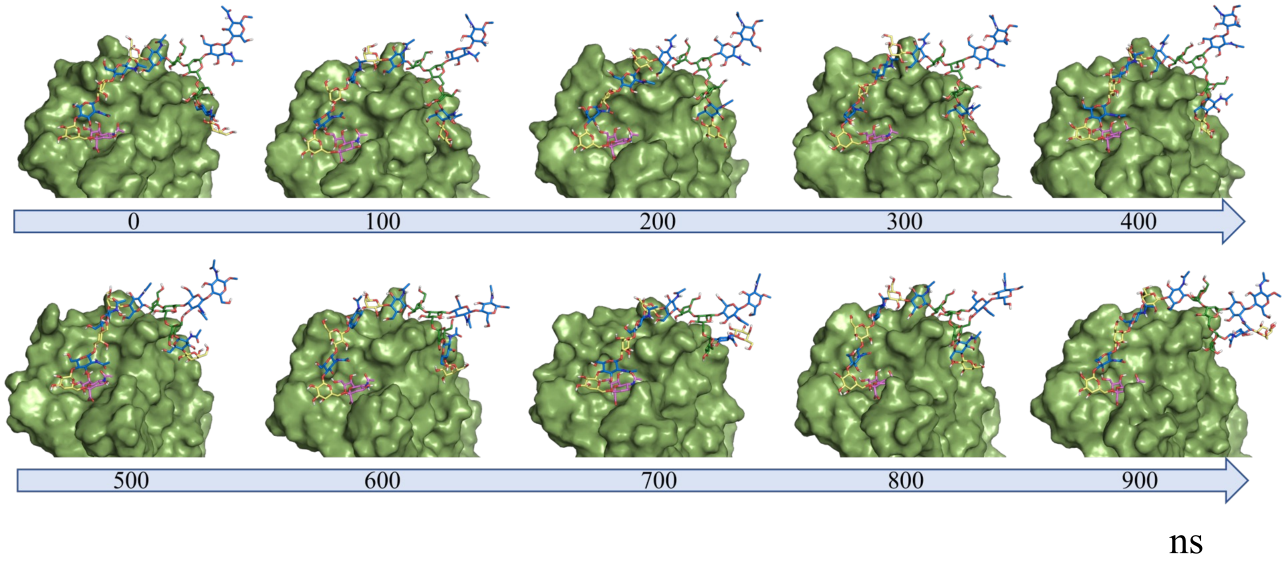

**8. Supplementary Fig. 10.** Snapshoot of the **2**/NL03 MD trajectory.

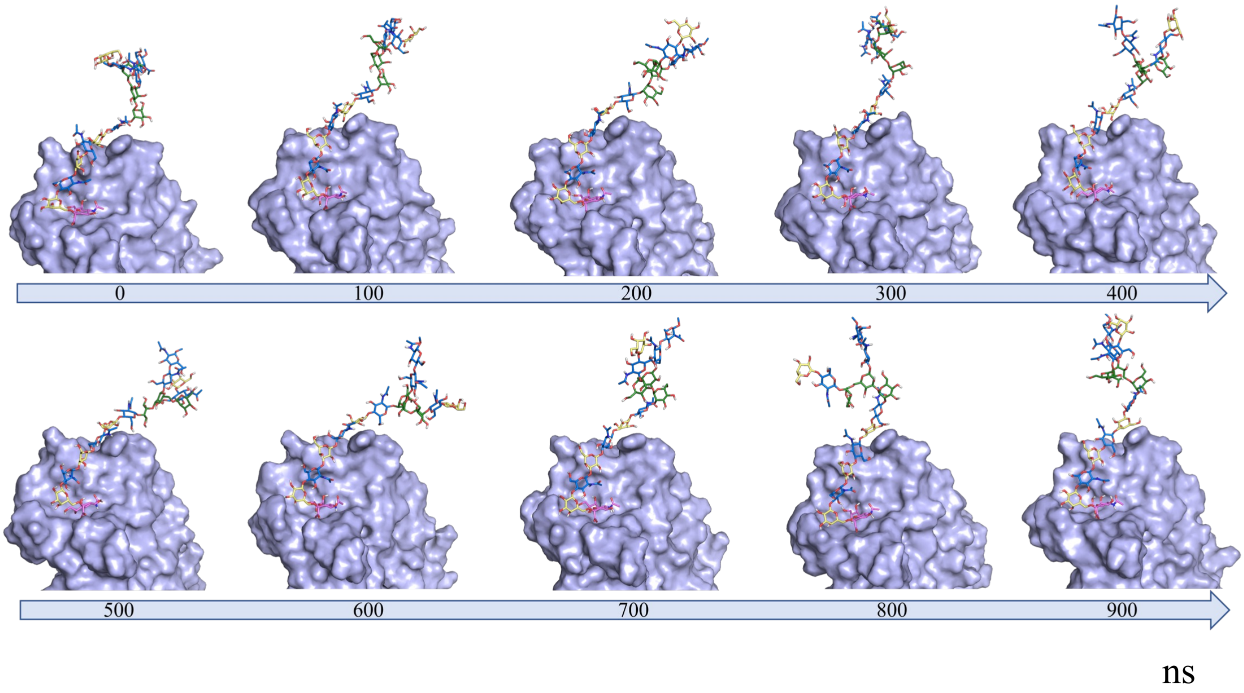
